# Mechanical forces and ligand-binding modulate *Pseudomonas aeruginosa* PilY1 mechanosensitive protein

**DOI:** 10.1101/2023.08.03.551776

**Authors:** Francisco J. Cao-Garcia, Jane E. Walker, Stephanie Board, Alvaro Alonso-Caballero

## Abstract

Bacteria initiate colonization and biofilm formation in response to mechanical cues caused by surface proximity. The protein PilY1 has been proposed as a key actor mediating mechanosensing. PilY1 is a calcium and integrin-binding protein with additional roles in host adhesion and functional regulation of the type IV pili (T4P), the appendages involved in twitching motility, and various aspects of the surface-associated life of bacteria. Due to its extracellular location and involvement in several surface processes, PilY1 is exposed to mechanical forces that could modulate its different roles. Herein, we explore the effect of mechanical forces and ligand binding on the conformational dynamics of the PilY1 C-terminal domain. Our single-molecule approach demonstrates that PilY1 acts as a ligand-modulated force sensor. At high forces, PilY1 unfolding occurs through a hierarchical sequence of intermediates. When calcium is bound to its cognate site linked to T4P regulation, there is a long-range mechanical stabilization affecting several PilY1 domains, which ensures the structural integrity of the protein. In the low-force regime, the integrin-binding domain of PilY1 exhibits calcium-tuned force sensitivity and conformational dynamics akin to those of mechanosensor proteins. Integrin binding to this domain occurs under force, inducing a shortening of its unfolded extension. Our findings suggest that the roles of the PilY1 C-terminal domain are force and ligand-modulated, which could entail a mechanical-based compartmentalization of its functions.

## Introduction

Mechanical forces play key roles in cell fate, triggering crucial events such as differentiation, proliferation, or motility^1^. Mechanical sensing and the responses elicited by force are well-known phenomena in eukaryotic cells; however, our understanding of how prokaryotes sense and respond to mechanical cues is still scarce^2^.

Although explored to a lower extent, it is recognized that mechanical stimuli promote critical life changes in prokaryotes^3–5^. Concretely, the mechanical cues induced by surface proximity stimulate the switch from a planktonic to a sessile lifestyle^6^. In pathogenic bacteria like *Pseudomonas aeruginosa*, surface detection is followed by colonization, biofilm formation, and virulence development towards a host^7^. It has been proposed that the protein PilY1 may act as a mechanosensor mediating surface detection^8^. This protein is located at the tip of the type IV pili^9^ (T4P), the motor-driven appendages that power bacterial translocation on substrates^3,10,11^. In addition to surface sensing, PilY1 is involved in T4P biogenesis and participates in several of its functions^12^, such as in twitching motility^13^, adhesion^14^, biofilm formation, and virulence^15,16^. Therefore, PilY1 stands out as a multipurpose protein with a pivotal role in numerous processes that govern the surface-associated life of bacteria^17^.

The different roles of PilY1 have been pinned down to specific features found in its sequence, which has enabled the compartmentalization of the protein into two functional sections (**Fig. 1A**). On the N-terminus, the von Willebrand A domain (vWA) has been assigned the mechanosensing role of PilY1. This function was inferred from the phenotype manifested after vWA domain deletion, which placed planktonic bacteria in a constitutive virulence state, a behavior typically observed after surface engagement^15^. The absence of the vWA domain mimicked the signaling triggered after PilY1 interaction with a surface, suggesting a role in mechanotransduction. More recently, it was found that the vWA domain could also have an adhesive function linked to conserved cysteine residues of its sequence. Mutation of these residues reduced the adhesion strength of bacteria and dwindled the responses characteristic of surface-committed cells^18^. While mounting evidence supports the involvement of the vWA domain in mechanosensing^19^, less is known about how force could regulate the functions of the C-terminal section of PilY1.

**Figure 1.**
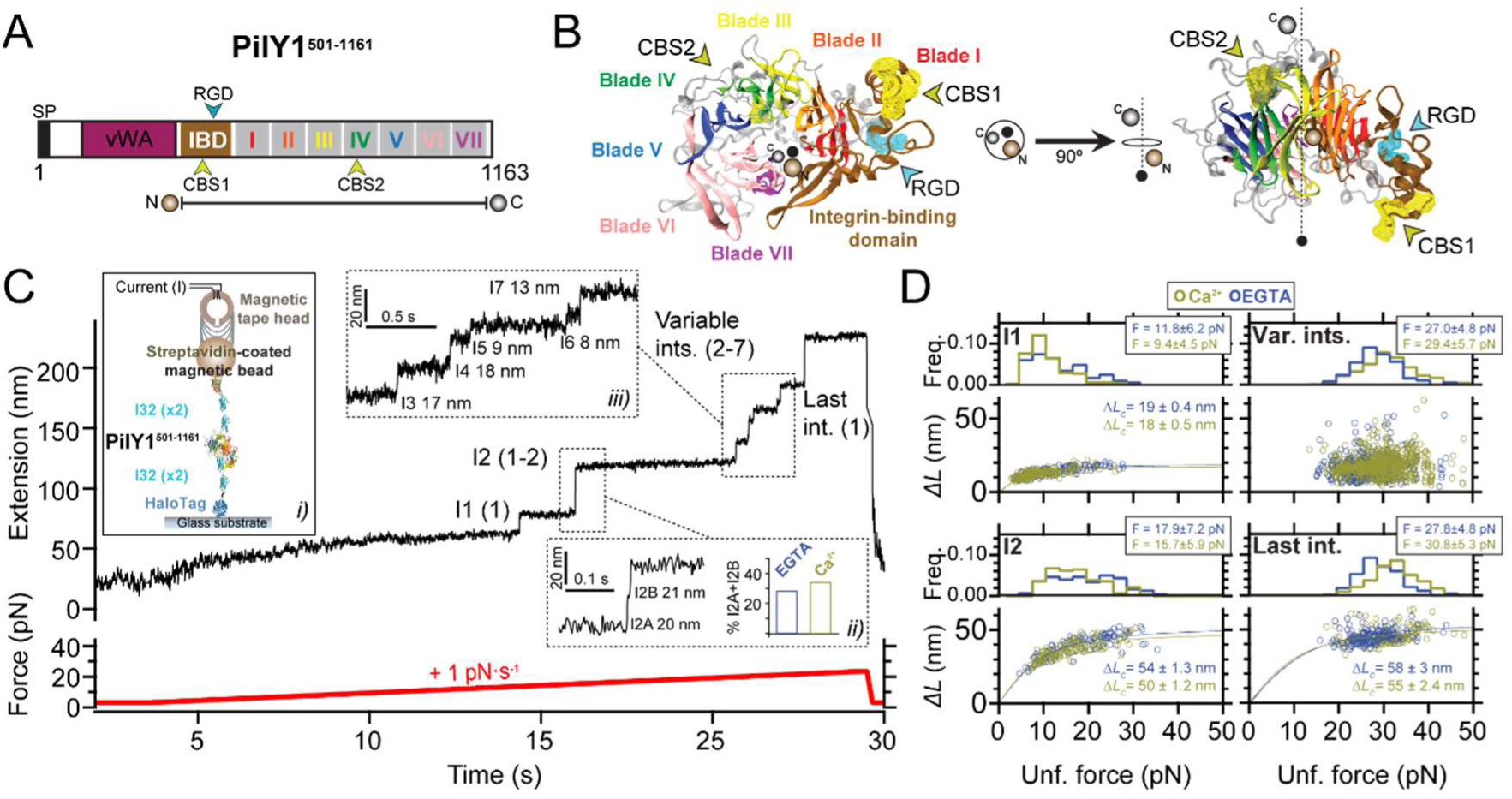
PilY1^501–1161^ mechanical hierarchy and Ca^2+^dual effect. **A)** Scheme of PilY1 protein sequence and structural features (SP: signal peptide; vWA: von Willebrand A domain; IBD: integrin-binding domain; RGD: integrin-binding site; CBS1: Ca^2+^-binding site 1; blades I to VII of the β-propeller; CBS2: Ca^2+^-binding site 2). The sequence employed spans from residue 501 to 1161. **B)** Crystal structure of PilY1^501–1161^ (prediction from Alphafold^36,37^, based on PDB:3HX6 and color assignation based on reference^13^) from top (left) and side (right) views. **C)** Magnetic tweezers force-ramp trajectory of PilY1^501–1161^ (inset *i*, see main text and methods for a full description of the chimeric protein construct and tethering) under a 1 pN·s^−1^ loading rate. PilY1^501–1161^ unfolds through multiple intermediates, which are classified based on their order of appearance: I1 (1 event), I2 (1 or 2 consecutive events, inset *ii*), variable intermediates (2-7 events of different sizes, inset *iii*), and last intermediate (1 event). **D)** Unfolding force distributions (top, mean±SD) and force-dependency of unfolding extensions (bottom) per intermediate group and condition (Ca^2+^ or EGTA). The extension (*ΔL*) vs unfolding force dependency is described with the freely jointed chain model (FJC) for polymer elasticity (lines in the bottom panels are fits of this model). Kuhn length values, *l_K_* in EGTA for I1, I2, and last intermediate are: 1.5 ± 0.1 nm, 1.0 ± 0.1 nm, and 0.7 ± 0.2 nm, and in Ca^2+^ are: 1.7 ± 0.1 nm, 1.1 ± 0.1 nm, and 0.9 ± 0.2 nm.

Unlike the vWA domain, the sequence of the C-terminal part of PilY1 is well-conserved across species, and its structure has been experimentally determined^13^. In *P. aruginosa* strains, the sequence spanning residues ∼640-1150 adopts a 7-bladed-β-propeller configuration. This type of fold comprises several structural repeats, termed blades, arranged around a central axis^13,20^ (**Figure 1A** and **1B**). Blade IV contains an EF-hand Ca^2+^-binding motif that modulates the extension and retraction cycles of the T4P. In the metal-bound state, PilY1 inhibits the retraction activity of the T4P machinery, which can exert pulling forces ranging 30-100 pN^21,22^. After Ca^2+^ release, pilus retraction is restored, indicating that the PilY1 C-terminal domain acts as a Ca^2+^-dependent switch that modulates T4P dynamics in biogenesis and twitching motility^13^. Further scrutiny of the PilY1 sequence identified an adhesin function between the vWA domain and β-propeller. PilY1 proteins from *P. aeruginosa* strains harbor an RGD motif that binds to the extracellular domains of integrin. A second Ca^2+^-binding site was found in the vicinity of the RGD motif, and it was demonstrated that integrin binding required both the RGD motif and Ca^2+^-binding on this site^23^. Integrins are transmembrane proteins present in the respiratory epithelium, a common niche of *P. aeruginosa* pathogenic strains^24,25^.

PilY1 plays a key role in surface-related processes and is exposed to mechanical perturbations. Understanding the force response and ligand-binding modulation of PilY1 could shed light on the molecular mechanisms underlying surface colonization. Herein, we have employed a single-molecule approach to address the nanomechanics of the PilY1 C-terminal section, which holds the ligand-binding sites responsible for T4P dynamics and host cell adhesion. Our results indicate that PilY1 is a mechanosensor protein that exhibits different behavior depending on the force load. In the high-force regime, PilY1 unfolds through multiple intermediates in a complex yet hierarchical sequence affected by Ca^2+^. When this metal is bound to its furthermost C-terminal site, which regulates T4P dynamics, PilY1 stability increases and becomes less sensitive to mechanical perturbations. In the low-force regime, the integrin binding domain undergoes folding and unfolding transitions in a 2 pN range, showing a steep force dependency characteristic of mechanosensitive proteins. Experiments with integrin enabled us to detect single binding events, which partially blocked the unfolding of its cognate domain and induced a shortening of the protein. We conclude that PilY1 C-terminal section exhibits a dual force dependency where its functions as an adhesin and T4P regulator could be invoked depending on the mechanical load experienced by bacteria during substrate colonization. Our work establishes a mechanical-based connection between the structure and functions of PilY1 and suggests that this bacterial protein has a force-sensing activity that extends beyond its N-terminal domain. These findings underpin the intricate and functional wealth of PilY1.

## Results

### Hierarchical unfolding and Ca^2+-^dependent mechanical stability

To explore the nanomechanics of PilY1, we employed single-molecule magnetic tweezers^26^. This technique allows the application of well-calibrated forces to single proteins and monitoring their conformational changes with high spatial, temporal, and force resolutions^27,28^. To conduct single protein measurements, we designed a chimeric polyprotein with features that enable its end-to-end specific tethering between a functionalized glass cover slide and a superparamagnetic bead, the force probe. Our protein of interest is the C-terminal section of PilY1 encompassing residues 501-1161 from the *P. aeruginosa* PAO1 strain (PilY1^501–1161^) (**Fig. 1A** and **1B**). In the chimeric construct (**Fig. 1C**, inset *i*), the N-terminal presents a HaloTag protein for covalent anchoring on the glass surface^29^, and the PilY1^501–1161^ sequence is flanked on both sides by two copies of the human titin I32 domain, which act as rigid molecular handles providing spacing between the surface and the bead. The C-terminal of the polyprotein contains a biotinylated AviTag sequence that allows the tethering to a streptavidin-coated superparamagnetic bead.

Our first goal was to investigate the mechanical stability and fingerprint of PilY1^501–1161^ under a steadily increasing force load. We also aimed to assess whether Ca^2+^ impacts its stability, as reported in protein mechanics^30–32^. For this reason, we conducted experiments with Ca^2+^ or the calcium chelator EGTA. **Fig. 1C** shows an unfolding trajectory of this construct, which was exposed to a loading rate of 1 pN·s^−1^ from its folded state up to its complete unfolding. As the load increases, the protein is stretched, and discrete jumps in the extension can be observed corresponding with the unfolding of PilY1^501–1161^ through multiple intermediates. These unfolding events are heterogeneous in extension and number; nevertheless, we noticed a conserved sequence. We assigned a number to each intermediate based on their order of appearance. In this trajectory, the least stable intermediate, I1, unfolds in a single 14 nm event at 13 pN. At 15 pN, we detect the unfolding of I2, which usually occurs as a single event, but, as shown in **Fig. 1C** inset *ii*, it can proceed through two sub-intermediates. Following I1 and I2, a population of unfolding events of heterogeneous quantity and size appears (**Fig. 1C** inset *iii*). The number of intermediates oscillates inter and intramolecularly (between unfolding pulses) and ranges from two large events to up to seven smaller extensions. Because of the difficulty of identifying reproducible conformations, we pooled them into a single group termed variable intermediates. The final ∼40 nm event in this trajectory, occurring at 22 pN, was named the last intermediate. Therefore, the unfolding pattern of PilY1^501–1161^ displays a clear fingerprint of four classes of events occurring hierarchically, with a conserved sequence of I1 (1 event), I2 (1-2 events), variable intermediates (2-7 events), and last intermediate (1 event).

In **Fig. 1D**, we show the unfolding force distribution (top) and the force-dependent step size (bottom) of each of the classes of intermediates detected in the presence of Ca^2+^ or EGTA. By collecting data from multiple trajectories and molecules, we obtain the scatter plots shown in **Fig. 1D**, which depict single unfolding events of each class occurring over a distribution of forces. The step size (*ΔL*) force-dependency of each class can be described by polymer models such as the freely jointed chain (FJC)^33^. From the fits of the data to this model, we can obtain the extension of the unfolded intermediates (*ΔL_c_*). In the case of I1, I2, and the last intermediate classes, the data points are well described by a single fit in both Ca^2+^ and EGTA, indicating that these events can be attributed to unique structural features in PilY1^501–1161^. In ∼30% of the trajectories, the I2 intermediate proceeds through two consecutive sub-intermediates (**Fig. 1C**, inset *ii*, and **Supplementary Fig. 1A**), which can be described with two separate fits whose combined *ΔL_c_* matches the *ΔL_c_* of I2 when detected as a single event (**Supplementary Fig. 1B**). In the variable intermediates class, the unfolding events cannot be described with an FJC fit due to its heterogeneity. Nevertheless, insights can be obtained from the unfolding force distributions of each group and the classification standard employed (**Fig. 1D**, top graphs). Ca^2+^ decreases the average unfolding force of both I1 and I2 (11.8±6.2 pN vs. 9.4±4.5 pN, for I1; 17.9±7.2 pN vs. 15.7±5.9 pN, for I2. mean±SD), also observed when I2 occurs in two events (**Supplementary Fig. 1B**). In contrast, Ca^2+^ increases the collective stability of the variable intermediates (27.0±4.8 pN vs. 29.4±5.7 pN) and the last intermediate (27.8±4.8 pN vs. 30.8±5.3 pN). Ca^2+^ has opposite effects on the PilY1^501–1161^ structure, lowering the mechanical stability of I1 and I2 while increasing that of the variable intermediates and last intermediate. Furthermore, and independent of the buffer condition, there is a gap in the unfolding forces between I1 and I2 (<25 pN), and the variable intermediates and the last intermediate (>25 pN).

PilY1^501–1161^ contains two Ca^2+^-binding sites, but from these observations, we cannot locate which group of intermediates could have these motifs. We sought to explore the variable intermediates group further, attending exclusively to their unfolding order sequence for their classification of mechanical stabilities. **Supplementary Fig. 1C** (color coding of the intermediates is unrelated to the colors chosen for **Fig. 1A** and **1B**) and **Supplementary Table S1** show the unfolding force distributions and summarize the results. In both EGTA and Ca^2+^, the first variable intermediate, I3, shows no differences in its average unfolding force (26.5 ± 5.0 pN vs. 25.8 ± 5.7 pN). In contrast, Ca^2+^ increases the unfolding forces of the following intermediates, from the second variable intermediate, I4, to the last intermediate. Comparisons inside each buffer condition reveal that in EGTA (**Supplementary Fig. 1C, Supplementary Table 1** and **2**), there is no significant change in the average unfolding force from I3 to the last intermediate (26.5 ± 5.0 pN vs. 27.8 ± 4.8 pN), suggesting that these structures have similar stabilities. With Ca^2+^ (**Supplementary Fig. 1C, Supplementary Table 1** and **3**), there is a significant ∼5 pN increase in the average mechanical stability between I3 and I4 (25.8 ± 5.7 pN vs. 30.4 ± 5.3 pN). After I4 and until the last intermediate, the distributions show no significant differences, indicating similar stability. These results suggest that the variable intermediate I4 is a plausible candidate to harbor one of the Ca^2+^-binding sites, and that the mechanical stabilization shift observed in subsequent intermediates could be due to Ca^2+^-binding on this intermediate.

### Ca^2+^ binding alters the unfolding pathway

The force-ramp experiments aided us in establishing a benchmark for further testing the nanomechanics of PilY1^501–1161^. We defined a clear mechanical signature of the protein, which consists of a hierarchical unfolding sequence of intermediates whose stabilities are affected by Ca^2+^. We next explored PilY1^501–1161^ unfolding dynamics under constant force. In these measurements, we can obtain a clearer picture of the force-dependent extensions of each of the intermediates, narrowing the variability observed in the force-ramp experiments. Since forces above 20 pN trigger the complete stretching of PilY1^501–1161^, we explored its unfolding dynamics from 20 to 40 pN to capture all the intermediates.

**Fig. 2A** shows a trajectory of PilY1^501–1161^ subjected to unfolding and refolding cycles at constant forces. Changing the force from 4 to 20 pN drives the protein from its folded state to its complete unfolding through multiple intermediates. After, the force is quenched to 4 pN to allow protein folding, which reaches the same folded extension. A subsequent 20 pN pulse reproduces the same pattern of unfolding intermediates. We identified the intermediate groups previously categorized (**Supplementary** Figs. 2A and 3A). In EGTA or Ca^2+^, the average number of variable intermediates is approximately 4. The number of total intermediates oscillates around 7-8, with a slightly higher count when Ca^2+^ is present. (**Fig. 2B**). The proportion of I2 unfolding events occurring in two steps doubled in the presence of Ca^2+^ (**Fig. 2A**, inset *iii*), explaining this increase. Fitting the FJC model to the average extensions of I1, I2, and the last intermediate yield similar results to those obtained in force ramp (**Supplementary Fig. 2D** and **3D**). However, the variable intermediates’ heterogeneity persists, as seen in **Fig. 2A** insets *i* and *ii*. For instance, I3 appears as a ∼18 nm extension in the first unfolding pulse, while the second shows a ∼6 nm step. This change in pattern affects all the variable intermediates except for I6, which shows a similar extension. This variability was detected intra and intermolecularly and cannot be explained by the reshuffling of the unfolding order of the intermediates between pulses, as their sizes differ. To rule out that the variable intermediates were not properly folding between pulses, we measured in all trajectories the extension of all the unfolding events in PilY1^501–1161^ and the extension of the unfolding events of the variable intermediates. **Fig. 2C** shows FJC fits to the combined extension of all the unfolding events (**), which yields an *ΔL_c_* ∼190 nm with both EGTA and Ca^2+^, and the variable intermediates (*), which collectively produce an *ΔL_c_* ∼70 nm in both conditions. These results indicate that despite the different number of events and extensions, the amount of polypeptide sequence released after unfolding is the same in all trajectories; therefore, these variable intermediates achieve a compact folded state.

**Figure 2.**
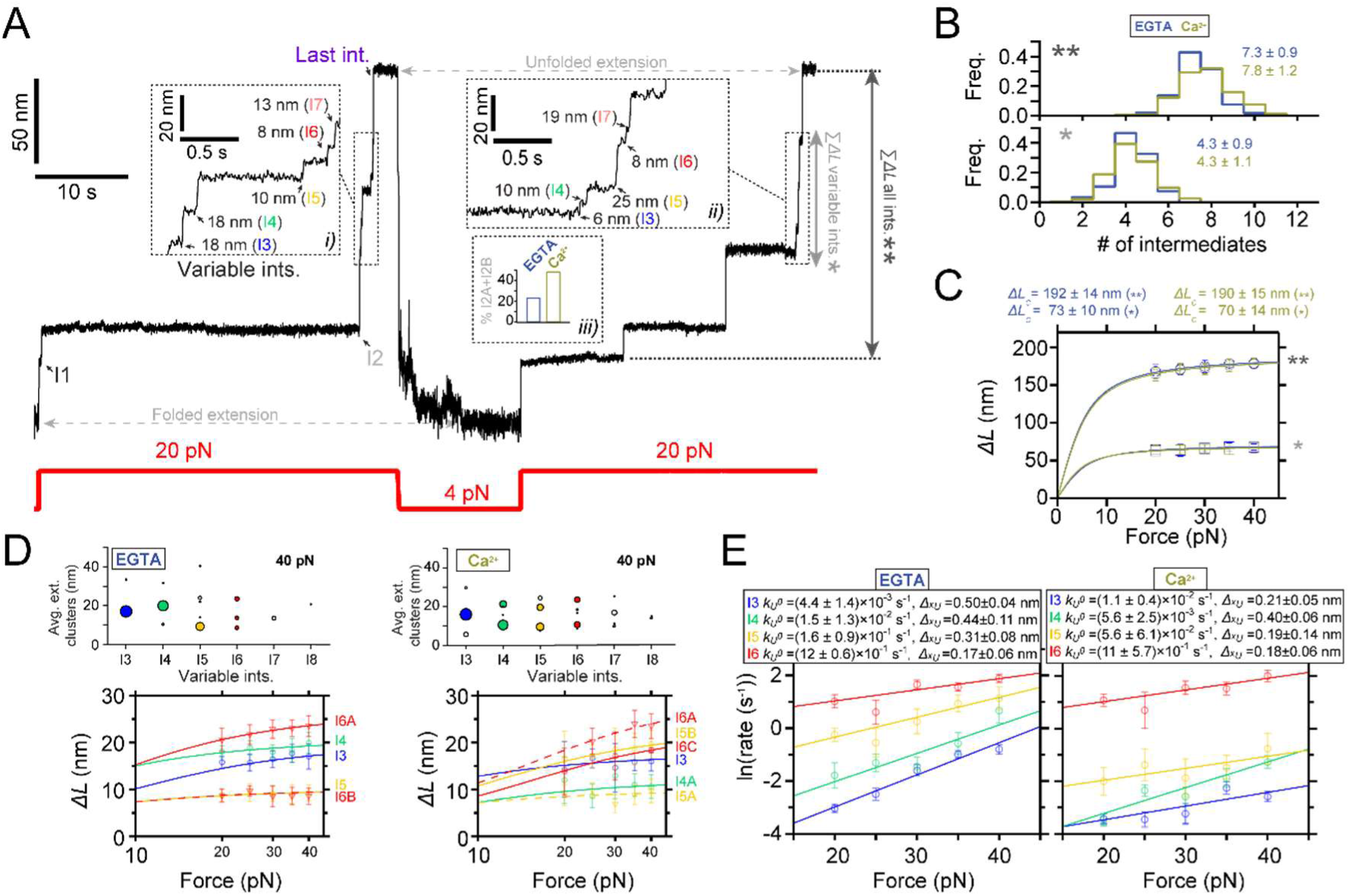
Ca^2+^-induced conformational heterogeneity and stabilization of PilY1 ^501–1161^. **A)** Magnetic tweezers trajectory of PilY1^501–1161^ exposed to unfolding and refolding cycles at different constant forces. From 4 pN, the force is jumped to 20 pN, and the sequential unfolding of PilY1^501–1161^ through multiple intermediates is detected (I1→I2→Variable intermediates→Last intermediate). The force is then quenched to 4 pN to allow protein folding, and later the 20 pN unfolding pulse is repeated, obtaining the same populations of intermediates. Insets *i* and *ii* highlight the conformational heterogeneity of the variable intermediates between consecutive unfolding pulses. In Ca^2+^, the proportion of I2 domains unfolding in two consecutive sub-intermediates doubled compared to EGTA (inset *iii*). Vertical arrows show the total length of the unfolding intermediates (**) and the variable intermediates (*). **B)** Distribution of the total number of unfolding intermediates in PilY1^501–1161^ (top, **) and variable intermediates (bottom, *) observed across trajectories in Ca^2+^ or EGTA. **C)** Force-dependent extension change of the combined extension of all PilY1^501–1161^ intermediates (**) and variable intermediates (*). Lines are fits of the FJC model to the average extension (mean±SD), and the resulting values of contour length increment (*ΔL_c_*) are shown above the graph for each condition. **D)** Cluster analysis results of the extension of the variable intermediates per order of unfolding and force in EGTA (left) or Ca^2+^ (right). On the top is plotted the average extension of the events classified inside each cluster identified for each intermediate at 40 pN. The size of the data points is proportional to the number of events in that cluster (most represented ones are colored). Below are shown the force-dependent extension changes (mean±SD) of the most populated conformations for each intermediate in EGTA (left) and Ca^2+^ (right). Solid and dashed lines are fits of the FJC model to the most populated and second most populated conformations, respectively (see **Supplementary Figure 2C and 3C**). The force axis is plotted on a logarithmic scale. **E)** Unfolding kinetics under force of the variable intermediates in EGTA and Ca^2+^. Each intermediate’s unfolding kinetics (mean±SEM) is fitted using Bel’s model for bond lifetimes. Above each plot is shown the value of the parameters determined: unfolding force at zero force (*k_U_^0^*) and distance to the transition state (*Δx_U_*).

Although the variable intermediates’ heterogeneity was prominent, a sequence of events repeated more frequently. To explore the existence of a canonical unfolding pathway, we resorted to clustering analysis, which enabled us to classify the variable intermediates based on their extension. This helped us to sort conformations, identify the most common ones, and resolve structural identities^34^ (top graphs in **Fig. 2D** and **Supplementary Figs. 2B** and **3B**). **Fig. 2D** bottom plots show FJC fits to the average extension of the most common intermediates in EGTA and Ca^2+^ (**Supplementary Fig. 2C** and **3C**). In EGTA, most events can be assigned to unique conformations and are well-described by individual fits. The I6 intermediate deviates from this trend with two alternative conformations, although one predominates (I6A). In contrast, the behavior with Ca^2+^ differs. Beyond the I3 intermediate, the upcoming intermediates show alternative conformations that occur in a similar frequency. For instance, I4 has a predominant configuration (I4A *ΔL_c_* ∼12 nm) that differs from the I4 population seen in EGTA (*ΔL_c_* ∼21 nm). In I5, the most represented conformation differs as well (I5B *ΔL_c_* ∼23 nm vs. I5A *ΔL_c_* ∼10 nm). For I6, the behavior differs as well in both conditions. These results suggest that Ca^2+^ alters the unfolding pathway of PilY1^501–1161^, changing the conformations of the intermediates after the variable intermediate I3.

We questioned next whether Ca^2+^ alters PilY1^501–1161^ unfolding kinetics. In **Fig. 2E**, we show the force-dependent unfolding kinetics for each intermediate. We used Bell’s approximation for bond lifetimes under force to model their unfolding kinetics^35^. From fits of the data to this model, we can obtain an extrapolation to zero force of the unfolding rates of each intermediate (*k_U_*^0^), and the distance to the transition state (*Δx_U_*). The comparison between conditions indicates that Ca^2+^ stabilizes intermediates I4 and I5, slowing their unfolding rates ∼2.7 and ∼2.9 times, respectively. Ca^2+^ has the opposite effect on I3, accelerating its unfolding ∼2.5 times, while I6 remains unchanged in both conditions. There is a pronounced change in the *Δx_U_* of I3 and I5, which drop substantially compared to the EGTA condition. This indicates that Ca^2+^ stabilizes PilY1^501–1161^, which confirms the observations made in force-ramp experiments.

### PilY1^501–1161^ harbors a mechanosensing-like structure

The previous sections described PilY1^501–1161^ nanomechanics at force regimes that promote its complete unfolding. The sequential unfolding of PilY1^501–1161^ indicates that I1 and I2 are less stable than the rest of the intermediates. Therefore, we conducted constant force measurements at loads below 10 pN, to test whether PilY1^501–1161^ exhibits dynamics in this force regime.

**Fig. 3A** shows a trajectory of PilY1^501–1161^ exposed to high and low mechanical loads. After unfolding at high force, the load is quenched to 5.5 pN, which promotes the collapse of the extended polypeptide and enables folding. At this force, PilY1^501–1161^ dwells over time across three levels of extension, which correspond with structures folding and unfolding. The lowest extension level corresponds with the folding of I1, hence the complete folding of PilY1^501–1161^ (**Fig. 3A** inset *i*). The second and third levels correspond with I1 unfolded extension (and I2 folded extension) and I2 unfolded level, respectively. In **Fig. 3A** inset *ii*, we show long dynamics of PilY1^501–1161^ at forces ranging from 4.5 to 6.0 pN, from where it can be seen how the increase in the mechanical load tilts the equilibrium towards the extended (unfolded state) levels of I1 and I2. Measuring at this force range allowed us to determine the force-dependent step size of I1 and I2 below 20 pN (**Supplementary Fig. 2D** and **3D**), which enabled us to assign their identity to these dynamics. From these long equilibrium dynamics (**Fig. 3A** inset *ii*), we determined the folding probability as a function of the force for I1 and I2 with EGTA or Ca^2+^ (**Fig. 3B**). As the mechanical load rises, the folded fraction of both intermediates decreases, adopting a sigmoidal shape. The force value at which the intermediates spend the same amount of time in the folded and unfolded state is defined as *P_50_*. Ca^2+^ shifts *P_50_* to lower values for I1 (4.7 to 4.4 pN) but increases it for I2 (5.7 to 5.9 pN). Moreover, Ca^2+^ changes the force dependency of I2 folding probability, which is broadened and spans 3 pN (from fully folded at 4 pN to unfolded at 7 pN), 0.5 pN more than with EGTA (4.0-6.5 pN).

**Figure 3.**
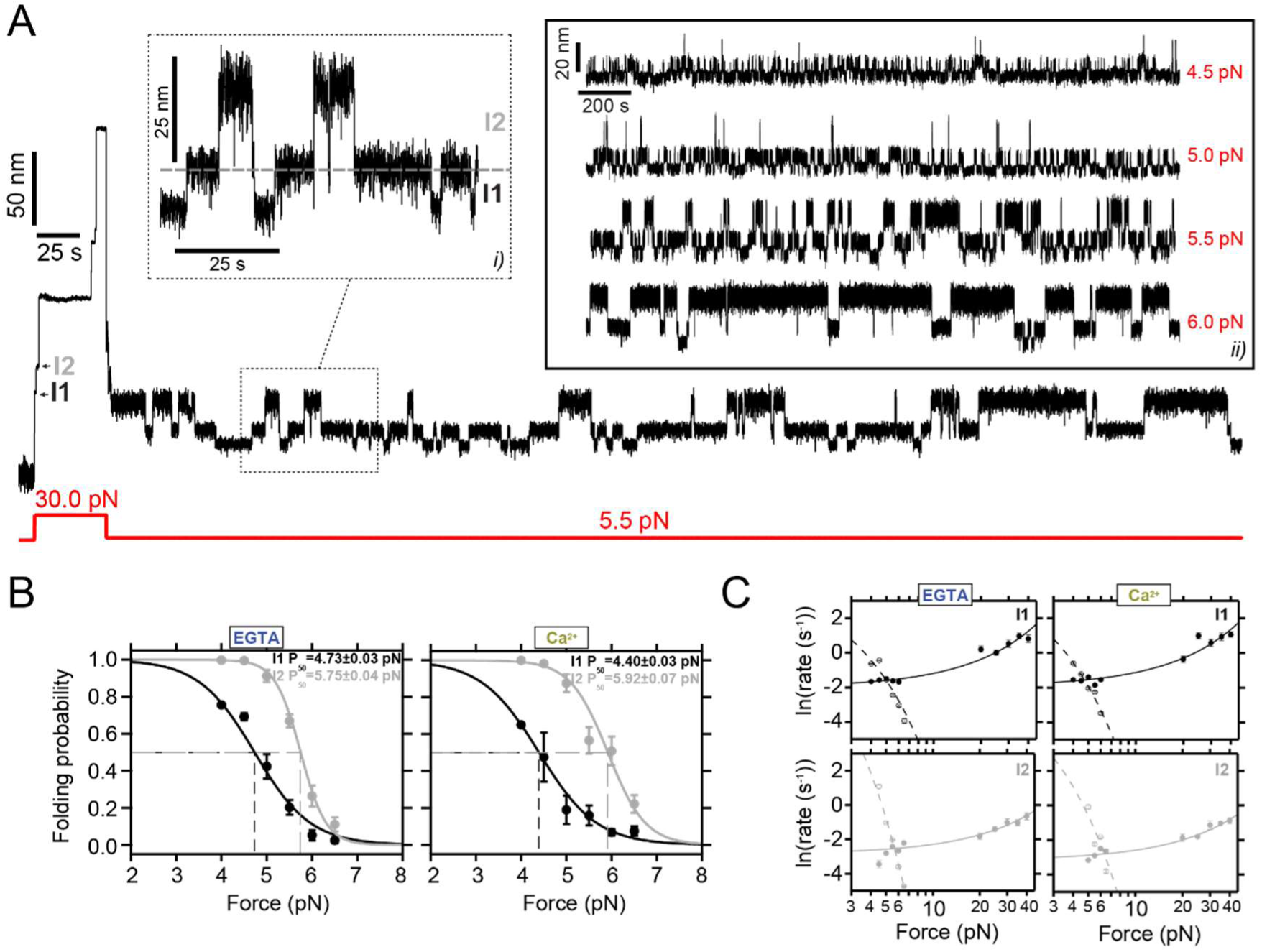
Conformational dynamics of PilY1^501–1161^ at low force reveal a mechanosensing-like behavior. **A)** Magnetic tweezers trajectory of PilY1^501–1161^ exposed to an unfolding pulse at 30 pN followed by a long 5.5 pN pulse. As shown before, the high-force pulse induces the unfolding of PilY1^501–1161^ through multiple intermediates. At low force, most of the protein refolds; however, part of the protein dwells across 3-levels of extensions (inset *i*) corresponding with the folded extension of I1 (lowest), the unfolded I1/folded I2 extension (middle), and unfolded extension of I2 (highest). The unfolding hierarchy previously observed at high forces is maintained in this force regime for I1 and I2, which also applies to their folding. Inset *ii* shows ∼2000 s-long trajectories of the dynamics of I1/I2 under different forces. The increase in the mechanical load promotes the occupation of the extended levels. **B)** Folding probability of I1 and I2 between 4 and 6.5 pN in EGTA and Ca^2+^. The folded fraction of both intermediates sharply decreases with force, following a sigmoidal trend (line fits). The coexistence force (*P_50_*) of I1 shifts to lower values, while that of I2 increases in the presence of Ca^2+^. Ca^2+^ also broadens the force range at which I2 exhibits folding and unfolding dynamics (∼2.5 pN in EGTA and ∼3 pN in Ca^2+^). **C)** Folding and unfolding kinetics under force of I1 and I2 in EGTA and Ca^2+^. The logarithm of the rate (mean±SEM) is plotted against the force in logarithmic scaling. I1 and I2 unfolding and folding kinetics are described with Bell’s model (fit lines) in both conditions (**I1**, empty circles and dashed lines for folding: EGTA *k_F_*^0^ = (7.1 ± 0.7) ×10 s^−1^, *Δx_F_* = - 4.83 ± 0.09 nm; Ca^2+^ *k_F_*^0^ = (13.4 ± 2.5) ×10 s^−1^, *Δx_F_* = - 5.62 ± 0.16 nm. Solid circles and lines for unfolding: EGTA *k_U_*^0^ = (13.7 ± 0.4) ×10^−2^ s^−1^, *Δx_U_* =0.33 ± 0.01 nm; Ca^2+^ *k_U_*^0^ = (14.4 ± 0.4) ×10^−2^ s^−1^, *Δx_U_* =0.34± 0.01 nm; **I2** empty circles and dashed line for folding: EGTA *k_F_^0^* = (2.4 ± 0.6) ×10^5^ s^−1^, *Δx_F_* = - 10.82 ± 0.18 nm; Ca^2+^ *k_F_^0^* = (2.3 ± 1.4) ×10^3^ s^−1^, *Δx_F_* = - 6.85 ± 0.44 nm. Solid circles and lines for unfolding: EGTA *k_U_^0^* = (5.8 ± 0.2) ×10^−2^ s^−1^, *Δx_U_* =0.23 ± 0.01 nm, Ca^2+^ *k_U_^0^* = (4.2 ± 0.2) ×10^−2^ s^−1^, *Δx_U_* =0.24 ± 0.01 nm)

We also tested if the kinetics of I1 and I2 were affected by Ca^2+^. **Fig. 3C** shows both intermediates’ folding and unfolding kinetics in EGTA or Ca^2+^. Using Bel’s model, we observe that the folding and unfolding kinetics of both I1 and I2 reproduce the observations from the folding probability. From the extrapolations to zero force, in Ca^2+^ I1 unfolds faster (∼1 time) and folds slower (∼1.9 times) than in EGTA, which explains the shifted folding probability. I2 folding rate at zero force is slowed down in Ca^2+^ by ∼100 times, which comes from a steep force-dependency change (EGTA *Δx_F_* = - 10.82 ± 0.18 nm vs. Ca^2+^ *Δx_F_* = - 6.85 ± 0.44 nm), while its unfolding rate is slowed down ∼1.4 times in comparison to the EGTA condition.

PilY1^501–1161^ dynamics at low force reveal that the hierarchy is conserved for I1 and I2, and both exhibit a Ca^2+^-tuned mechanical sensitivity in a narrow range of forces, which showcases the prototypical behavior of mechanosensors. These results indicate that at least one of them contains one Ca^2+^-binding site, although both are affected by metal binding to this motif.

### The integrin-binding domain is the mechanosensitive structure of PilY1^501–1161^

After demonstrating the mechanosensing-like behavior of PilY1^501–1161^, we wanted to link the observed intermediates with structural modules of this protein. PilY1^501–1161^ sequence contains the β-propeller fold and a stretch of 141 residues in the N-terminus that contains the RGD and Ca^2+^-binding motifs responsible for integrin binding. Unlike the β-propeller, the structure of this section from PilY1 has not been resolved (**Fig. 1B** shows a prediction from AlphaFold^36,37^).

To elucidate which of the intermediates of PilY1^501–1161^ corresponds to this sequence containing the RGD and first Ca^2+^-binding motifs, we synthesized a second construct that only contains the β-propeller, named PilY1^642–1161^ (**Fig. 4A**). **Fig. 4B** shows a trajectory of PilY1^642–1161^ exposed to unfolding and refolding pulses. The unfolding trajectories show no sign of I1 and I2, but only the variable intermediates and last intermediate groups (**Fig. 4B**, and **Supplementary Fig. 4A** and **5A**). The average number of total intermediates drops to ∼5, while the number of variable intermediates remains unchanged (**Fig. 4C**), which indicates the loss of two domains. The complete unfolded extension of all intermediates (**Fig. 4D**) is ∼60 nm shorter than PilY1^501–1161^ (**Fig. 2C**), but the variable intermediates complete extension is similar. After cluster analysis (**Supplementary Fig. 4B** and **5B**), the variable intermediates reproduce the conformational structures observed in PilY1^501–1161^ (**Fig. 4E** and **F**, **Supplementary Fig. 4C** and **5C**), and the effect that Ca^2+^ has in both the unfolding pathway and kinetics (**Fig. 4G**, and **Supplementary Table 4**). Therefore, the second Ca^2+^-binding site, located in blade IV of the β-propeller, is responsible for mechanically stabilizing the variable intermediates. Together with the results obtained in **Figure 2**, the intermediate I4 seems the most plausible candidate for being blade IV in the β-propeller.

**Figure 4.**
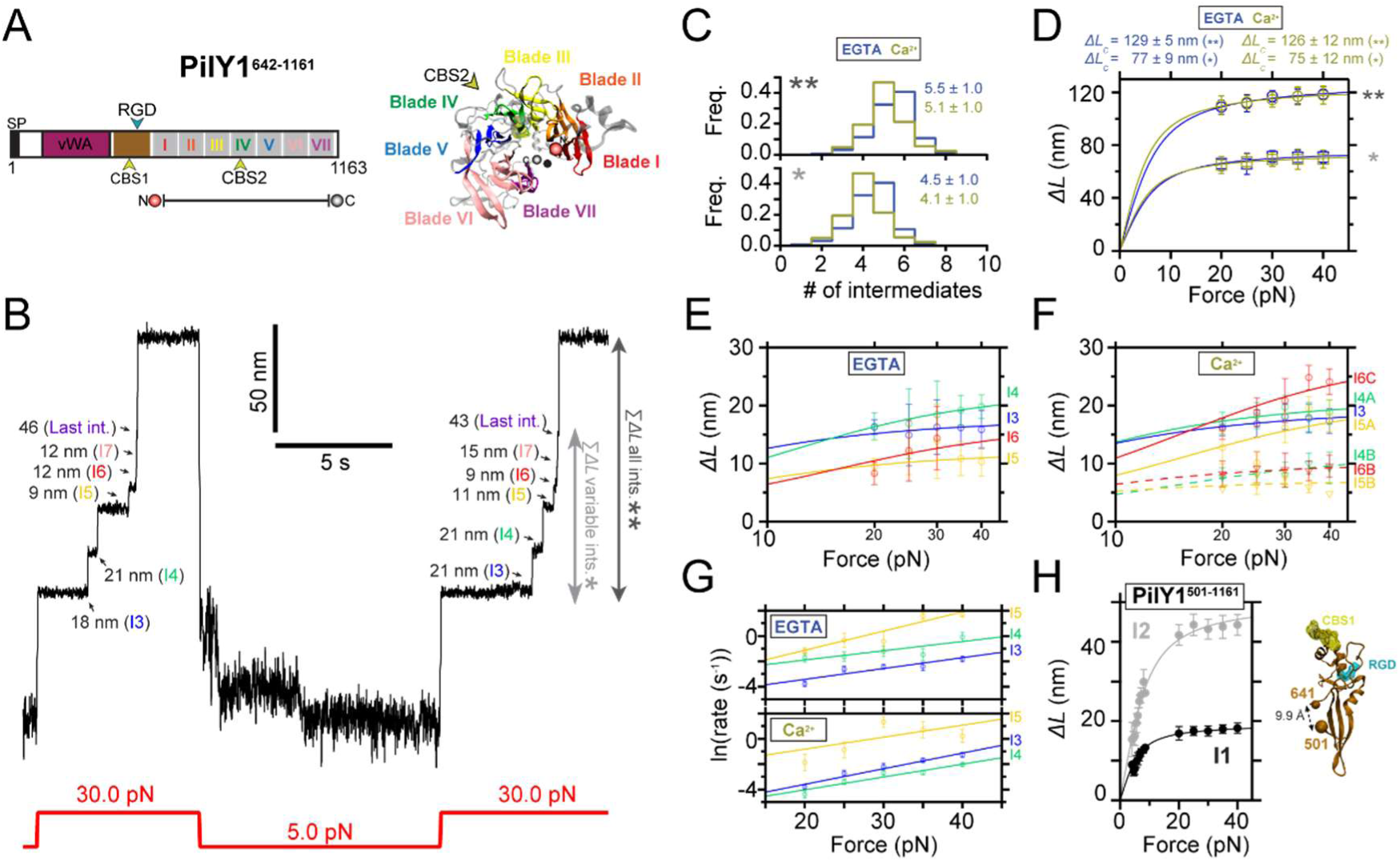
Nanomechanics of PilY1^642–1161^ indicates that the integrin-binding domain is responsible for the mechanosensor-like behavior observed at low force. **A)** Scheme of PilY1 protein and crystal structure (PDB:3HX6^13^) of the sequence employed spanning residues 642 to 1161. **B)** Magnetic tweezers trajectory of PilY1^642–1161^ exposed to unfolding and folding cycles. Removing the sequence 501-641 yields a mechanical fingerprint identical to PilY1^501–1161^ but lacking the intermediates I1 and I2, where we only detect the variable intermediates and last intermediate groups. **C)** Distribution of the total number of unfolding intermediates in PilY1^642–1161^ (top, **) and variable intermediates (bottom, *) observed across trajectories in Ca^2+^ or EGTA. **D)** Force-dependent change in the combined extension of all PilY1^642–1161^ intermediates (**) and variable intermediates (*). Lines are fits of the FJC model to the average extension (mean±SD), and the resulting values of contour length increment (*ΔL_c_*) are shown above the graph for each condition. **E and F)** Force-dependent extensions (mean±SD) of the most populated conformations determined with cluster analysis for each intermediate in EGTA (**E**) and Ca^2+^ (**F**). Solid and dashed lines are fits of the FJC model to the most populated and second most populated conformations, respectively (see **Supplementary Fig. 4C** and **5C**). The force axis is plotted on a logarithmic scale. **G)** Unfolding kinetics under force of the variable intermediates in EGTA and Ca^2+^. Each intermediate’s unfolding kinetics (mean±SEM) is fitted using Bel’s model for bond lifetimes (see **Table S4**). **H)** Based on its sequence length and predicted structure, the integrin-binding domain could be the I2 intermediate observed in PilY1^501–1161^, which yields an *ΔL_c_* ∼50 nm (plot on the left). By contrast, I1 would not be a good candidate to hold the integrin-binding site due to its smaller extension.

These findings demonstrate that the sequence spanning residues 501 to 641 is involved in the dynamics observed at 4.0-6.5 pN. However, the removal of this 140-long residue sequence has a larger than expected impact on the total length of the protein. The predicted structure (**Fig. 4H**) indicates that the N and C-termini of this integrin-binding domain are close to each other (*L_0_* ∼1 nm). Assuming a length of 0.36 nm·residue^−1^, the 140-long polypeptide would yield a *ΔL_c_* ∼ 49 nm, in agreement with the values obtained for I2 (**Fig. 4H** and **Supplementary Fig. 2D** and **3D**). While these observations support the connection between the 501-641 sequence and I2, the missing ∼60 nm length of PilY1^641–1161^ accounts for the loss of I2 and I1, which was never detected. Measurements of the complete extension of the PilY1^501–1161^ and PilY1^642–1161^ constructs from 0.5 pN to its full unfolding at 40 pN, reveal that both proteins achieve their full predicted extensions based on their sequence length (**Supplementary Fig. 6**). Hence, the absence of I1 signature in PilY1^642–1161^ could be due to its inability to fold properly if the integrin-binding domain is not present.

### Integrin binding blocks the unfolding of I2

The evidence from the previous section suggests that the intermediate I2 could be the putative structure that contains the RGD and Ca^2+^-binding site responsible for integrin binding. To prove this hypothesis, we conducted binding experiments in the presence of the extracellular domains of human integrin αVβ5 (**Fig. 5A**), which was shown the integrin with the highest affinity to PilY1^23^. To conduct these experiments, we used a buffer supplemented with calcium, magnesium, and manganese (binding buffer), since these metals are important for the structural integrity of integrin^23,38^. Before proceeding with the binding experiments, we characterized PilY1^501–1161^ nanomechanics in this new buffer. **Supplementary Figures 7** and **8** summarize these results in force-ramp and constant force measurements, respectively. The protein exhibits a mixed mechanical behavior, where the unfolding force distributions of I1 and I2 are like the ones observed with EGTA, but the variable and last intermediates reproduce the Ca^2+^ condition results (**Supplementary Fig. 7**). PilY1 can bind with similar affinity Ca^2+^ and Mn^2+^, but not magnesium^23^; hence, Mn^2+^ could be interfering with Ca^2+^ binding in the integrin-binding domain. By contrast, the unfolding force distributions of the intermediates modulated by the second Ca^2+^-binding site are identical to the Ca^2+^ buffer condition, which suggests that both sites have different affinity for Mn^2+^.

**Figure 5.**
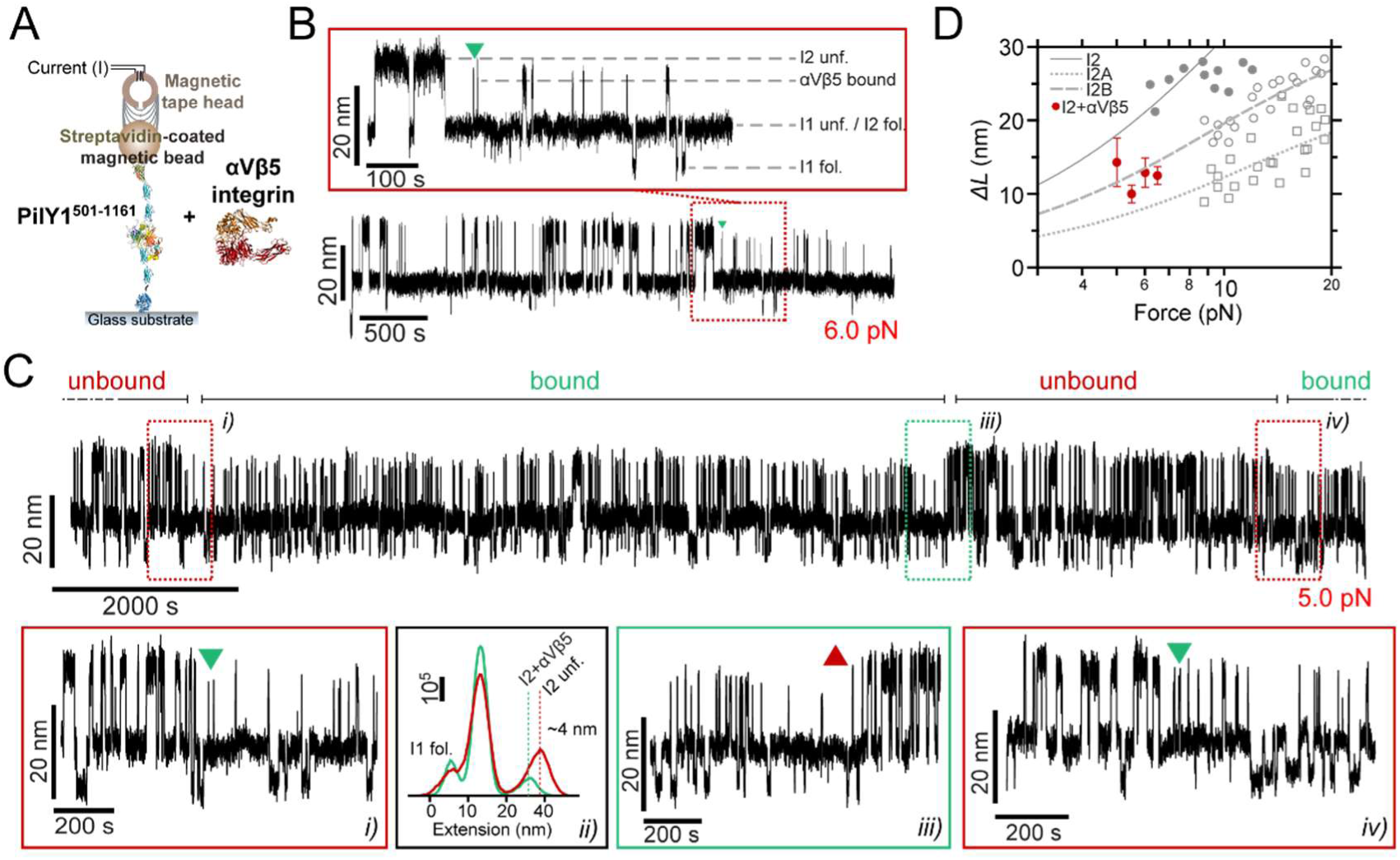
Integrin binds to I2 in PilY1^501–1161^ under force and prevents its full unfolding. **A)** Diagram of the magnetic tweezers binding experiment with PilY1^501–1161^ and the extracellular domains of the human integrin αVβ5 heterodimer. **B)** Magnetic tweezers trajectory of PilY1^501–1161^ held at 6 pN in the presence of integrin. I1 and I2 domains fold and unfold in equilibrium, and integrin binding is detected as a shortening of the unfolded extension of I2 (inset). **C)** ∼4-hour-long trajectory at 5 pN where integrin binds (insets *i* and *iv*) and unbinds (inset *iii*) to PilY1^501–1161^ I2 domain, yielding a ∼4 nm change in the extension. Inset *ii* shows the extension distribution of the protein before binding (red) and after binding (green). Bar shows the number of points. **D)** Comparison of the force-dependent extension changes of the integrin-bound I2 (mean±SD) with force-ramp data from non-bound I2 and its two sub-intermediates I2A and I2B. Fitting the data with the FJC model reveals that integrin-bound I2 force-dependent extension falls within the fit for I2B sub-intermediate, which indicates that integrin-binding prevents the unfolding of I2A.

After setting the basis of PilY1^501–1161^ nanomechanics in binding buffer, we conducted integrin binding experiments under force. **Fig. 5B** details the fingerprint for integrin binding events. At 6 pN, I1 and I2 domains fold and unfold in equilibrium visiting the three different levels of extension. As shown in the inset, at the moment indicated with a green arrow, the protein stops visiting the unfolded extension of I2, but the dynamics continue unchanged below this level. In **Fig. 5C**, we show a trajectory of PilY1^501–1161^ held at 5 pN for a long period of time in the presence of integrin. Insets *i* and *iv* indicate binding events and inset *iii* one unbinding event. In the bound state the I2 domain cannot reach its unfolded extension, but a shorter one. After unbinding, I2 recovers its corresponding extension at the measured force (inset *ii* shows the protein extension distribution before and after binding). We measured the average extension of this new conformation of I2 across the forces where integrin binding was detected (**Fig. 5D**). When compared with the force-dependent extension of naive I2, we can see that integrin-bound I2 average extension falls within the FJC fit of the sub-intermediate I2B. This would indicate that integrin binding partially blocks the unfolding of I2, and that the sub-intermediate I2A cannot extend under this condition. Hence, I2 is the conformational structure holding the integrin-binding domain that spans from residue 501 to 641.

## Discussion

Bacteria colonize and adhere to substrates upon recognition of the mechanical stimuli elicited by surface proximity^8,19^. These cues trigger signaling pathways in free-swimming cells that promote their attachment to target substrates and the formation of biofilms. This change of lifestyle reprograms gene expression and, in pathogenic bacteria, kickstarts the deployment of virulence factors, having important implications for infection progression^24,39^.

The ability of bacteria to detect mechanical forces has been linked to their motor-driven appendages, the flagella and the T4P^10,40^. The T4P mediate twitching motility and are essential for the surface-attached life of bacteria. PilY1 plays a crucial part in T4P biogenesis and functions, while also serving as a mechanosensor and adhesive protein. Due to its location at the tip of the T4P, PilY1 carries out its functions while exposed to mechanical stress. Here, we have explored how ligand-binding and mechanical forces impact PilY1 dynamics.

The sequence comprising the residues 501-1161 of PilY1 contains ligand-binding motifs that control T4P dynamics and host adhesion. Most of this sequence adopts a 7-bladed-β-propeller fold, where blade IV contains the Ca^2+^-binding site that regulates T4P extension and retraction cycles. Forces above 20 pN lead to the sequential hierarchical unfolding of the domains of this heteropolyprotein. Under force, a polyprotein comprised of independent domains of varying levels of stability will usually undergo the unfolding of the least stable domains first before the more stable ones^41^. In the PilY1 constructs studied (**Figures 1-5**, and **Supplementary Figures 2-5**), we observe a tightly conserved unzipping pattern where the unfolding of each domain depends on the unfolding of the previous one in the sequence, which reminds a Matryoshka doll-like configuration. This suggests the existence of long-range contacts and allosteric interactions over the entire structure of the protein^42–44^. The apparently conserved unfolding pathway is challenged by the heterogeneous conformations exhibited by the variable intermediates group. Ca^2+^ binding to its cognate site in blade IV increases the mechanical stability of the variable intermediate I4 and delays the unfolding of the subsequent domains (**Figures 2** and **4**, **Supplementary Figures 1** and **7**). Furthermore, this metal increases the frequency of conformations sparsely visited in EGTA (**Figures 2** and **4**, and **Supplementary Figures 2** to **5**), indicating that metal binding impacts the mechanical properties and the unfolding pathway of the protein^30,45–48^. The fact that the total unfolding extension of the variable intermediates is always the same confirms that this region of PilY1 folds between unfolding cycles (**Figures 2** and **4**). A source of conformational variability could be that, after quenching the force and allow folding, the variable intermediates experience misfolding events where their contacts are not achieved intradomain, but instead non-native interactions are established with neighboring modules^49^. If that was the case, the unfolding rates of the variable intermediates could not be explained as single processes (**Figures 2** and **4**), but as a multitude of kinetic processes, each of them departing from a different folded state. Hence, the hierarchical pattern observed for I1, I2, and last intermediate groups also applies inside the variable intermediates sequence.

Our findings indicate that PilY1 β-propeller has a partially conserved unfolding pathway that Ca^2+^ alters when bound to blade IV. A rough estimation of the expected unfolded extension of each blade of the β-propeller and its candidate intermediates is shown in **Supplementary** Figure 9. PilY1 is not a canonical β-propeller fold^20^, and blades V-VII have a different number of strands than blades I-IV, which are four-stranded antiparallel β-sheets. However, the estimated contour length of all blades is similar, and assigning their identities is not straightforward. Furthermore, blades V and VI share one strand^13^, which makes it even more challenging to devise what kind of unfolding intermediate or intermediates could generate. Ultimately, the core population of the variable intermediates could be narrowed down to four different structures of *ΔL_c_* ∼10-20 nm (I3 to I6), and the alternative conformations observed can be explained as the rupture of substructures spanning different blades.

From a biological standpoint, the stabilizing effect of Ca^2+^ in the β-propeller could be relevant in T4P dynamics regulation. *P. aeruginosa* T4P exerts pulling forces ∼30 pN that overlap with the force-ranges we have tested, and where Ca^2+^ has shown its stabilizing effect^22^. How the Ca^2+^-bound state of PilY1 antagonizes T4P retraction is not yet clear, but one study hypothesized that PilY1, in association with other pilus-tip proteins involved in T4P biogenesis, might block the full retraction of the T4P beyond the outer membrane PilQ secretin pore, acting as a cork^9^. In this scenario, Ca^2+^-induced mechanical stabilization of PilY1 would maintain the bulky structure of the β-propeller and prevent its unfolding, blocking further pilus retraction. This would ensure the structural integrity of the pilus-tip complex for priming pilus polymerization in a subsequent extension cycle. PilY1 orthologs exhibit different Ca^2+^ affinity and its effect in T4P dynamics vary from *P. aeruginosa*, which highlights that Ca^2+^-mediated binding and T4P regulation could be finely tuned to the chemical environment of bacteria^50–52^.

PilY1 mechanical behavior at high force can match with its expected function as a T4P retraction inhibitor, but its dynamics in the low force regime resemble a force-sensing activity linked to its adhesin function. The I2 intermediate seems the most plausible candidate to harbor the motifs responsible for integrin binding, not only by its dimensions but also for the signature conformation it adopts upon binding (**Figures 1, 3**, and **5**). Shortening of the unfolded extension of proteins after ligand binding has been reported in the proteins talin and vinculin, keystones of eukaryotic mechanotransduction^53,54^. Integrin binding to PilY1 prevents the full extension of the I2 domain but, unlike the talin-vinculin interaction, the remaining structure of I2 can still undergo folding and unfolding transitions. The conformation of this half-way structure matches the force-dependent extension of the sub-intermediate I2B; hence, integrin binding prevents the unfolding of I2A. These sub-intermediates unfold almost at the same time in a sequence I2A-I2B and are noticeable when pulling at high forces. Their frequency increases with Ca^2+^ (**Figure 2** and **Supplementary Fig. 8**), which suggests that metal-binding in I2 could promote a structural organization of the binding pocket that facilitates integrin recruitment^23^. Furthermore, Ca^2+^ binding in I2 seems to activate an allosteric mechanism that lowers the stability of I1, which might have implications for integrin binding modulation under tensile stress. This structural connection between I1 and I2 seems plausible since the elimination of I2 abrogates the mechanical signature of I1 (**Figure 4**), pointing out that both domains are interdependent for folding. This force and postranslational modulation of the nanomechanics of the integrin-binding domain and I1 could entail an evolutionary tuning to the mechanical environment of the respiratory epithelium, as it has been proposed for other bacterial adhesive structures^55,56^. Integrin binding has been shown to occur in the absence of force^23^; nevertheless, the allosteric interactions that seem to govern PilY1 mechanics could play a significant role in vivo, promoting or hindering host adhesion depending on the mechanical environment experienced by the bacterium.

Overall, our approach to PilY1 dynamics provides mechanistic details of how force could tune some of its functions at the nanoscale. We propose that PilY1 functions are mechanically compartmentalized; their recruitment is dependent on the stability of the structures that harbor them, ligand binding, and the mechanical load. Future work should address the nanomechanics of the vWA domain, given its center role in bacterial mechanostransduction, and the nanomechanics of the entire PilY1. Our findings hint at a complex network of interactions between structural modules, and PilY1 functions under force could depend on its structure as a whole.

## Methods

### Cloning and expression of chimeric constructs

All reagents employed were purchased from Sigma-Aldrich, unless otherwise indicated. The sequence of the proteins PilY1^501–1161^ and PilY1^642–1161^ from *P. aeruginosa* PAO1 strain were optimized for expression in *Escherichia coli*. Both sequences were ordered (GeneArt, Thermo Fisher Scientific) with a 5’-end Kpn2I and a 3’-end NheI restriction sites, which enabled their cloning into a modified pFN18A (HaloTag^®^) T7 Flexi^®^ Vector (Promega). The expression cassette contained, from 5’ to 3’-end, an N-terminal HaloTag protein, four copies of the human titin I32 domain, HisTag sequence, and AviTag sequence. The sequence between the second and third I32 domain gene copies contained the Kpn2I and NheI restriction sites, which allowed the insertion of the proteins of interest in sense. Cloning and DNA amplification procedures were carried out in *E. coli* TOP10 strain (Thermo Fisher Scientific), and sequence fidelity was confirmed (GeneWiz).

Protein expression was conducted in *E. coli* BL21 Star strain (Thermo Fisher Scientific). Cell culture was grown to OD_600_∼0.6 in LB broth containing 100 µg·mL^−1^ carbenicillin at 37°C and 250 rpm shaking, and then 1 mM IPTG (Invitrogen) was added to induce protein expression at 25°C for 3 hours and constant shaking. After, cells were pelleted (4°C, 4000 x *g*, 20 min), and resuspended in wash buffer (50 mM NaPi at pH 7.0, 300 mM NaCl, 20 mM imidazole, 10% v/v glycerol) supplemented with EDTA-free protease inhibitor (Roche). Cells were then incubated for 30 min on ice with 1 mg·mL^−1^ lysozyme, 5 µg·mL^−1^ DNase I, 5 µg·mL^−1^ RNase A, and 10 mM MgCl_2_. Then, the cells were mechanically lysed in a French press (G. Heinemann) and cell debris was pelleted (4°C, 40,000 x *g*, 1 h). The supernatant was incubated with HisPur Cobalt resin (Thermo Fisher Scientific) at 4°C for 1h on an orbital shaker. The His-tagged proteins were eluted from the resin with wash buffer containing 250 mM imidazole, and subsequently biotynilated in vitro at room temperature for 1 h. After, the protein solution was further purified by size exclusion chromatography in a Superdex^®^ 200 Increase 10/300 GL column (Cytiva). The protein was eluted from the column in storage buffer (10 mM Hepes pH 7.2, NaCl 150 mM, 1 mM EDTA, 10% v/v glycerol), aliquoted, snap frozen in liquid N_2_, and stored at −80°C until use.

### Single-molecule magnetic tweezers measurements

Single-molecule experiments were conducted on custom-built magnetic tweezers microscopes, either using a configuration consisting of a pair of permanent magnets, as previously described^27^, or a magnetic tape head^57^. In the magnetic tape head configuration, the force is applied to the magnetic bead-bound proteins with a magnetic tape head (Brush Industries), which is fixed on a custom-designed CNC aluminum piece (named card-reader). In the base of the card-reader lies the experimental fluid chamber where the protein under study is immobilized on the bottom glass (see below for details). The 150 µm thickness of the bottom glass places the sample-located surface at a 300 µm-distance from the 25 µm gap of the magnetic tape head, which establishes a known current-force relationship^58^. The 25 µm gap of the magnetic tape head is aligned with the optical path of the microscope. The base of the card-reader piece contains an opening to accommodate a 100X oil-immersion plan apochromat objective (Zeiss) that enables the visualization of the reference and magnetic beads located on the bottom glass of the fluid chamber. The objective position was controlled with a P-725 nanofocusing piezo actuator (Physike Instrumente), and the image was acquired with a CMOS camera (Ximea). Image processing was conducted with a custom-written program (C++/Qt) that allowed a ∼1.5 kHz image sampling frequency. This software communicated with a NI USB-6289 multifunction DAQ card (National Instruments) which enabled data acquisition and controlling the piezo position and the current supplied (hence, the force applied to the protein) to the magnetic tape head. The magnetic tape head electrical supply came from two 12 V lead-acid batteries and the current was kept under feedback with a custom-build PID controller. Applying currents in the 0-1000 mA range allowed the application of forces spanning from 0 to 42 pN using Dynabead M270 streptavidin-coated superparamagnetic beads (Invitrogen).

Fluid chamber preparation involved the cleaning and functionalization of a 40×24 mm bottom glass (150 µm thickness) and a 22×22 mm top glass (Ted Pella), as previously described^27^. After cleaning, the top glass was treated with repel-silane for 30 min and the bottom glass was silanized with a 1% v/v solution of 3-(aminopropyl)trimethoxysilane in ethanol for 10 min. After functionalization, both types of glasses were rinsed in 100% ethanol, dried with N_2_, and curated in an oven at 100°C for at least 1 h.

The fluid chambers were assembled by melting a bowtie-shaped parafilm spacer placed between the top glass on the bottom glass at 80°C for 5 min. A laser-cutter was used to generate the parafilm spacers shape. After sandwiching between glasses, the parafilm shape created two wells flanking the top glass that allowed buffer exchange during experiments. Further fluid chamber functionalization steps carried out at room temperature involved the sequential treatment with 1% v/v solution of glutaraldehyde in 0.1 M phosphate buffer (PBS) for 1 h, covalent immobilization of a 0.05% w/v solution of amino-functionalized polystyrene reference beads (2.5-2.9 µm in diameter, Spherotec) in PBS for 20 min, extensive rinse with PBS, and covalent anchoring of 20 µg·mL^−1^ HaloTag amine ligand (Promega) in PBS overnight. After HaloTag ligand functionalization the fluid chambers were passivated for at least 3 h with blocking buffer containing 1% w/v sulfydryl-blocked BSA (Lee Biosolutions) in 20 mM Tris-HCl pH 7.4, 150 mM NaCl, and 0.01% w/v NaN_3_.

To conduct force experiments, the fluid chambers were incubated with a ∼3 nM solution of the chimeric constructs (PilY1^501–1161^ and PilY1^642–1161^) at room temperature for 30 min. The unbound protein was removed by flowing buffer from one well of the fluid chamber and removing from the other. The basic composition of the buffer used in the experiments contained 10 mM Hepes pH 7.2, 150mM NaCl, and 10 mM ascorbic acid. Depending on the condition, this buffer contained either 2 mM EGTA, 2 mM CaCl_2_, or binding buffer (2 mM CaCl_2_, 1 mM MgCl_2_, and 1 mM MnCl_2_). The fluid chamber was then fixed with double-side tape (Tesa) to a custom-made metal stamped fork. The unit formed by the fork-fluid chamber assembly was placed in the base of the card-reader piece, leaving the fluid chamber below the magnetic tape head. The fork was then attached to an XY linear stage (Newport) which allowed to move the fluid chamber and scan different areas during the experiments.

The experiments started by flowing a 1:30 dilution of streptavidin-coated superparamagnetic beads, previously passivated with blocking buffer at 4°C, for >1 h, and constant rotation at 5 rpm. The beads and the surface-immobilized proteins were allowed to react for 2 min in the absence of force. Then, a constant force of 2-4 pN was applied to remove unspecific-bound beads, and the fluid chamber was further flowed with experimental buffer. From here, the different force protocols described in the manuscript were applied. For the integrin binding experiments, PilY1^501–1161^ proteins were held at forces that allowed folding and unfolding transitions of the I1 and I2 intermediates, and then a binding buffer solution containing 200 nM of recombinant αVβ5 human integrin extracellular domains (R&D Systems) was slowly added to one of the wells of the fluid chamber, while simultaneously removing the previous buffer volume from the other well until complete buffer substitution.

### Analysis

All the analysis was carried out with Igor Pro 8.0 software (Wavemetrics). Magnetic tweezers recordings were captured at ∼1.0-1.5 kHz framerate and trajectories were smoothed with a 151 box-sized 4^th^ order Savitzky-Golay filter. A two-sample Kolmogorov-Smirnov (K-S) test was conducted to test the differences in the unfolding force distribution of the intermediates in force-ramp. Step size determination of the unfolding intermediates was determined by measuring the extension before and after the transition, and for the equilibrium dynamics at low force a multi-gaussian fit was employed to find the center of the extension levels and measure their distance. The force-dependent extension of the intermediates was fitted to the FJC model for polymer elasticity^33^:

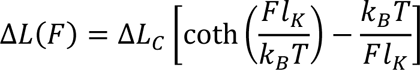

Where Δ*L*_C_ is the increment of contour length in nm, *l*_K_ is the Kuhn length in nm, and *k*_B_*T* is the thermal energy as 4.11 pN·nm. Behavior at high forces is dominated by the contour length, while Kuhn length is determined by low-force behavior. Hence, Kuhn length is only reported for those measurements and intermediates which exhibit folding and unfolding events in the low force regime (I1 and I2).

The force-dependent folding probability of the intermediates I1 and I2 was fitted with a sigmoidal function:

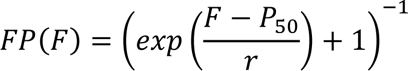

Where *P*_5O_ is the coexistence force (in pN) between the folded and unfolded state, and *r* is the rate in pN^−1^.

Dwell time analysis in the unfolding trajectories was measured from the moment the desired force is achieved (∼40 µs) until each of the intermediates unfold. Since their unfolding is sequential and requires that the previous structure unfolds first, their unfolding times were corrected based on this. The same procedure was used for the dynamics registered at low force. Due to their long nature a custom-written procedure was written to automatically detect the extension levels and assign the folding and unfolding dwell times of I1 and I2 intermediates. Folding and unfolding rates were calculated as the inverse of the average dwell times, and their force dependency was fitted with the Bell model^35^:

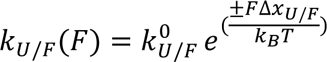

Where 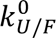 is the unfolding/folding rate extrapolation to zero force, and Δ*x_U/F_* is the distance to the transition state for unfolding and folding.

Cluster analysis of the variable intermediates was done with R Statistical Software (v 4.3.0; R Core Team 2023)^59^ using the libraries “factoextra”^60^ and “cluster”^61^. The k-means algorithm and the Euclidean distance were used to establish the differences in the extension of the conformations of the variable intermediates. We tested several k-means solutions varying the number of clusters from 1 to 20 for each force, protein construct, and buffer condition. We conducted the average silhouette analysis method to find the optimal number of clusters per condition, and this value was used to cluster the data.

## Acknowledgments

This project has received funding from the European Union’s Horizon 2020 research and innovation program under the Marie Sklodowska-Curie grant agreement No 101028879. This work was supported by and performed at the Francis Crick Institute, which receives its core funding from Cancer Research UK (CC0102), the UK Medical Research Council (CC0102), and the Wellcome Trust (CC0102). F. J. C-G. was supported by the Spanish Ministry of Economy and Competitiveness and the European Regional Development Fund (RTI2018-095802-B-I00). J. W. and S. B. were supported by the Leverhulme Trust (RL 2016-015). A. A-C. was funded by a European Commission Marie Sklodowska-Curie (MSCA-IF) fellowship (NIOBMT-101028879), and a Ramón y Cajal fellowship (RYC2021-031965-I) funded by Ministerio de Ciencia e Innovación (Spain) and the European Union (NextGeneration EU/PRTR). The authors thank Pablo Mateos-Gil, Marta Martín, and Rafael Rivilla for helpful discussions and comments

## Author contributions

A. A-C. conceived the project and designed research. S. B, J. W., and A. A-C. cloned and expressed proteins. A. A-C. conducted experiments and data analysis. F. J. C-G. conducted data analysis. A. A-C. drafted the paper, and all the authors contributed to revising and editing the manuscript.

