## Supplementary Information for "Mechanical forces and ligand-binding modulate *Pseudomonas aeruginosa* PilY1 mechanosensitive protein"

#### Supplementary Tables

**Table S1. Comparison of unfolding force distributions of PilY1 intermediates in EGTA and $Ca^{2+}$.** Significant differences between distributions are indicated with an asterisk (* p<0.05) in the last column, after the p-value of a two-sample Kolmogorov-Smirnov (K-S) test.

| Intermediate | Unfolding force, pN (mean±SD) EGTA | # events EGTA | Unfolding force, pN (mean±SD) $Ca^{2+}$ | # events $Ca^{2+}$ | K-S test, p-value (*p<0.05) |
|---|---|---|---|---|---|
| **I1** | 11.8±6.2 | 141 | 9.4±4.5 | 172 | $3 \times 10^{-4}$ (*) |
| **I2**<br>**I2A**<br>**I2B** | 17.9±7.2<br>19.5±5.9<br>19.6±5.9 | 101<br>40<br>40 | 15.7±5.9<br>16.8±5.7<br>16.8±4.7 | 113<br>59<br>59 | 0.02 (*)<br>0.04 (*)<br>0.05 |
| **Variable ints.** | 27.0±4.8 | 541 | 29.4±5.7 | 611 | $6 \times 10^{-13}$ (*) |
| **I3** | 26.5±5.0 | 140 | 25.8±5.7 | 172 | 0.30 |
| **I4** | 26.9±5.0 | 140 | 30.4±5.3 | 172 | $3 \times 10^{-7}$ (*) |
| **I5** | 27.0±4.6 | 132 | 30.9±5.1 | 157 | $9 \times 10^{-9}$ (*) |
| **I6** | 27.2±4.5 | 97 | 30.9±5.8 | 86 | $5 \times 10^{-5}$ (*) |
| **I7** | 27.5±5.2 | 27 | 32.2±5.7 | 19 | 0.014 (*) |
| **I8** | 32.0±6.7 | 5 | 32.1±3.0 | 3 | 0.7 |
| **I9** | N/A | N/A | 32.3 | 1 | N/A |
| **Last int.** | 27.8±4.8 | 141 | 30.8±5.3 | 172 | $3 \times 10^{-7}$ (*) |

**Table S2. Comparison of unfolding force distributions (values in Table S1) between intermediates in EGTA.** Significant differences between distributions are indicated with an asterisk (* p<0.05), after the p-value of a two-sample Kolmogorov-Smirnov (K-S) test. Cells in green highlight that between I3 and subsequent intermediates, there is no significant difference in their unfolding force distributions (see **Supplementary Table 1** and **Supplementary Figure 1C**)

| Int. | I1 | I2/I2A/I2B | I3 | I4 | I5 | I6 | I7 | I8 | Last int. |
|---|---|---|---|---|---|---|---|---|---|
| **I1** | | $2 \times 10^{-7}$ (*)<br>$2 \times 10^{-7}$ (*)<br>$2 \times 10^{-7}$ (*) | $1 \times 10^{-40}$ (*) | $3 \times 10^{-41}$ (*) | $3 \times 10^{-42}$ (*) | $3 \times 10^{-37}$ (*) | $2 \times 10^{-15}$ (*) | $2 \times 10^{-15}$ (*) | $1 \times 10^{-43}$ (*) |
| **I2**<br>**I2A**<br>**2B** | | | $2 \times 10^{-15}$ (*)<br>$2 \times 10^{-8}$ (*)<br>$2 \times 10^{-8}$ (*) | $9 \times 10^{-16}$ (*)<br>$4 \times 10^{-8}$ (*)<br>$4 \times 10^{-8}$ (*) | $1 \times 10^{-16}$ (*)<br>$2 \times 10^{-8}$ (*)<br>$2 \times 10^{-8}$ (*) | $5 \times 10^{-16}$ (*)<br>$2 \times 10^{-8}$ (*)<br>$2 \times 10^{-8}$ (*) | $6 \times 10^{-7}$ (*)<br>$5 \times 10^{-5}$ (*)<br>$5 \times 10^{-5}$ (*) | $9 \times 10^{-4}$ (*)<br>$9 \times 10^{-3}$ (*)<br>$9 \times 10^{-3}$ (*) | $1 \times 10^{-18}$ (*)<br>$2 \times 10^{-9}$ (*)<br>$2 \times 10^{-9}$ (*) |
| **I3** | | | | 0.9 | 0.8 | 0.5 | 0.8 | 0.07 | 0.2 |
| **I4** | | | | | 0.9992 | 0.93 | 0.998 | 0.1 | 0.6 |
| **I5** | | | | | | 0.995 | 0.98 | 0.1 | 0.8 |
| **I6** | | | | | | | 0.9997 | 0.1 | 0.98 |
| **I7** | | | | | | | | 0.2 | 0.9995 |
| **I8** | | | | | | | | | 0.2 |
| **Last int.** | | | | | | | | | |

**Table S3. Comparison of unfolding force distributions (values in Table S1) between intermediates in $Ca^{2+}$.** Significant differences between distributions are indicated with an asterisk (* $p<0.05$), after the p-value of a two-sample Kolmogorov-Smirnov (K-S) test. Cells in orange highlight that between I3 and subsequent intermediates, there is a significant difference in their unfolding force distributions. After, from I4 to the last intermediate, the distributions show no difference (see **Supplementary Table 1** and **Supplementary Figure 1C**).

| Intermed. | I1 | I2/I2A/I2B | I3 | I4 | I5 | I6 | I7 | I8 | Last int. |
|---|---|---|---|---|---|---|---|---|---|
| **I1** | | $6 \times 10^{-16}$ (*)<br>$8 \times 10^{-15}$ (*)<br>$8 \times 10^{-15}$ (*) | $6 \times 10^{-67}$ (*) | $6 \times 10^{-69}$ (*) | $6 \times 10^{-68}$ (*) | $7 \times 10^{-48}$ (*) | $3 \times 10^{-15}$ (*) | $1 \times 10^{-3}$ (*) | $3 \times 10^{-70}$ (*) |
| **I2<br>I2A<br>2B** | | | $1 \times 10^{-31}$ (*)<br>$3 \times 10^{-16}$ (*)<br>$3 \times 10^{-16}$ (*) | $8 \times 10^{-4}$ (*)<br>$1 \times 10^{-25}$ (*)<br>$5 \times 10^{-25}$ (*) | $6 \times 10^{-41}$ (*)<br>$3 \times 10^{-8}$ (*)<br>$1 \times 10^{-25}$ (*) | $3 \times 10^{-28}$ (*)<br>$4 \times 10^{-20}$ (*)<br>$2 \times 10^{-19}$ (*) | $4 \times 10^{-11}$ (*)<br>$5 \times 10^{-10}$ (*)<br>$5 \times 10^{-10}$ (*) | $2 \times 10^{-3}$ (*)<br>$3 \times 10^{-3}$ (*)<br>$3 \times 10^{-3}$ (*) | $2 \times 10^{-41}$ (*)<br>$2 \times 10^{-26}$ (*)<br>$9 \times 10^{-26}$ (*) |
| **I3** | | | | $2 \times 10^{-15}$ (*) | $1 \times 10^{-15}$ (*) | $3 \times 10^{-11}$ (*) | $2 \times 10^{-6}$ (*) | 0.02 (*) | $1 \times 10^{-16}$ (*) |
| **I4** | | | | | 0.9 | 0.99 | 0.2 | 0.6 | 0.93 |
| **I5** | | | | | | 0.9995 | 0.5 | 0.7 | 0.9999 |
| **I6** | | | | | | | 0.5 | 0.7 | 0.9999 |
| **I7** | | | | | | | | 0.98 | 0.4 |
| **I8** | | | | | | | | | 0.7 |
| **Last int.** | | | | | | | | | |

**Table S4. Summary of the data from the Bell model fits in the main text Figure 4G.**

| Intermed. | $k_U^0$ ($s^{-1}$) EGTA | $\Delta x$ (nm) EGTA | $k_U^0$ ($s^{-1}$) $Ca^{2+}$ | $\Delta x$ (nm) $Ca^{2+}$ |
|---|---|---|---|---|
| **I3** | $(5.9 \pm 2.7) \times 10^{-3}$ | $0.35 \pm 0.06$ | $(2.3 \pm 0.9) \times 10^{-3}$ | $0.51 \pm 0.05$ |
| **I4** | $(3.5 \pm 2.5) \times 10^{-2}$ | $0.30 \pm 0.10$ | $(2.3 \pm 1.0) \times 10^{-3}$ | $0.42 \pm 0.05$ |
| **I5** | $(1.6 \pm 0.9) \times 10^{-2}$ | $0.62 \pm 0.07$ | $(6.6 \pm 7.7) \times 10^{-2}$ | $0.39 \pm 0.15$ |
| **I6** | $(1.3 \pm 0.8) \times 10^{-1}$ | $0.43 \pm 0.08$ | $(9.4 \pm 1.1) \times 10^{-1}$ | $0.43 \pm 0.08$ |

### Supplementary Figures

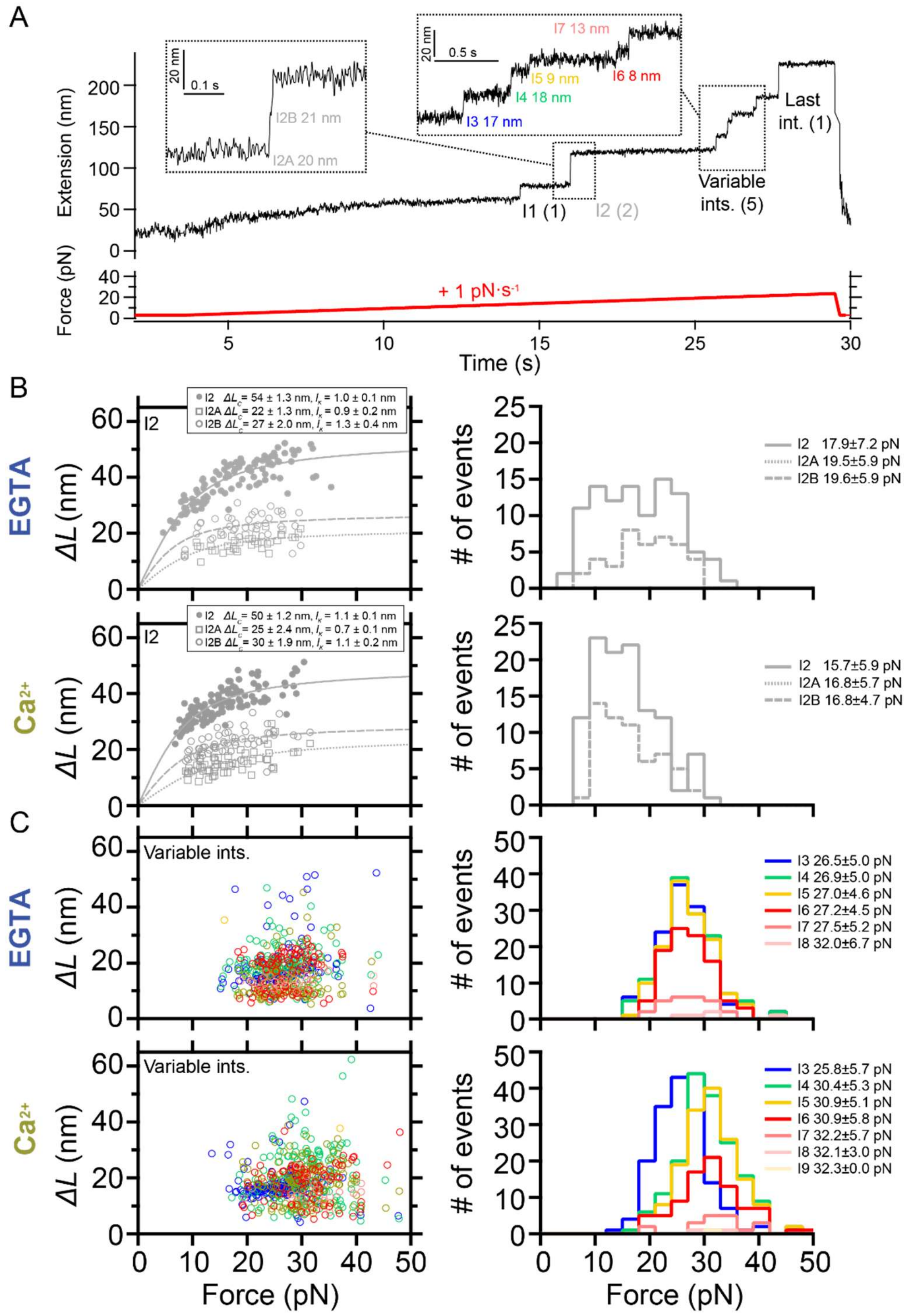


**Supplementary Figure 1. Mechanical characterization of PilY1$^{501\text{-}1161}$ in force-ramp (1 pN·s$^{-1}$) in the presence of $Ca^{2+}$ or EGTA.** **A)** Force-ramp trajectory of PilY1$^{501\text{-}1161}$ highlighting the unfolding of I2 through two sub-intermediates and the variable intermediates, which are numbered and colored based on their order of appearance in the unfolding sequence. **B)** On the left, force-dependent extensions of I2 and its two sub-intermediates and their calculated $\Delta L_c$ and $l_K$ from fits of the data with the FJC model in EGTA and $Ca^{2+}$ (top and bottom, respectively). On the right, unfolding force distributions (mean±SD) of I2 and its sub-intermediates in EGTA and $Ca^{2+}$ (top and bottom, respectively). **C)** On the left, force-dependent extensions of the variable intermediates (colored as in **A**) in EGTA and $Ca^{2+}$ (top and bottom, respectively). On the right, unfolding force distributions of the variable intermediates in EGTA and $Ca^{2+}$ (top and bottom, respectively). I4 and subsequent intermediates show an average unfolding force shifted to higher values with $Ca^{2+}$.

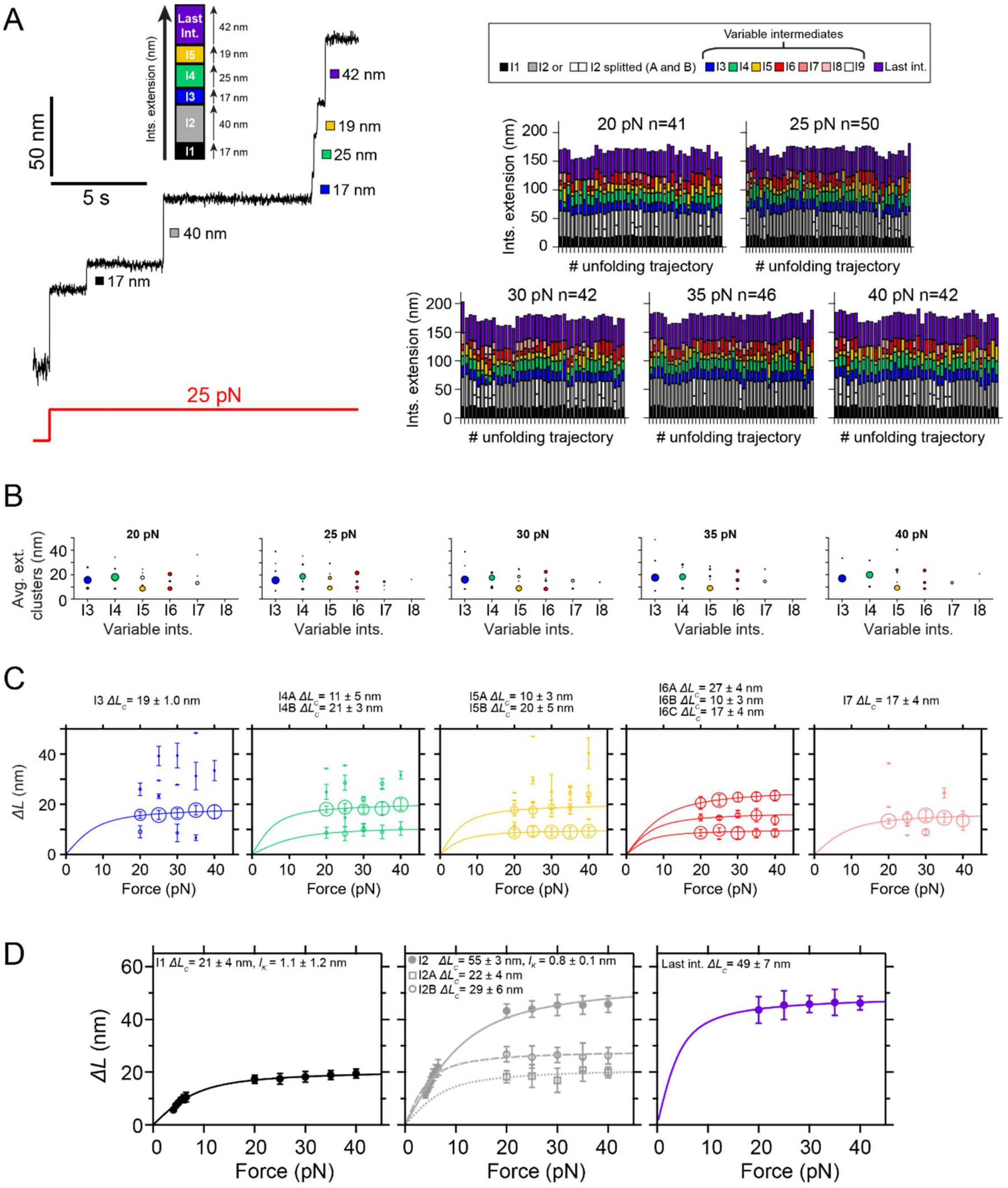


**Supplementary Figure 2. Mechanical characterization of PilY1$^{501\text{-}1161}$ in constant force with EGTA and identification of the most prevalent conformations of the variable intermediates.** Unfolding under constant force of PilY1$^{501\text{-}1161}$ in the presence of EGTA and classification of the intermediates by unfolding order and color. The category plots on the right show the stacked extensions of each intermediate per trajectory across the five unfolding forces tested (20, 25, 30, 35, and 40 pN). **B)** Cluster analysis of the extension of the variable intermediates per order of unfolding and force in EGTA. Each plot shows the average extension of the events classified inside each cluster and for each intermediate across the five tested forces. The size of the data points is proportional to the number of events in that cluster (most represented ones are colored). **C)** Force-dependent extensions (mean±SD) of the most populated conformations for each intermediate (I3 to I7). The size of the data points is proportional to the number of events in that cluster. When the extension of the clusters identified in **B** is represented for each intermediate across the five unfolding forces tested, it is possible to identify the different conformations of each intermediate and model the data with individual fits of the FJC model ($\Delta L_c$ data for each population shown above). **D)** Force-dependent extensions (mean±SD) of the I1, I2 (as a single and as two events), and last intermediates and their fitting with the FJC model.

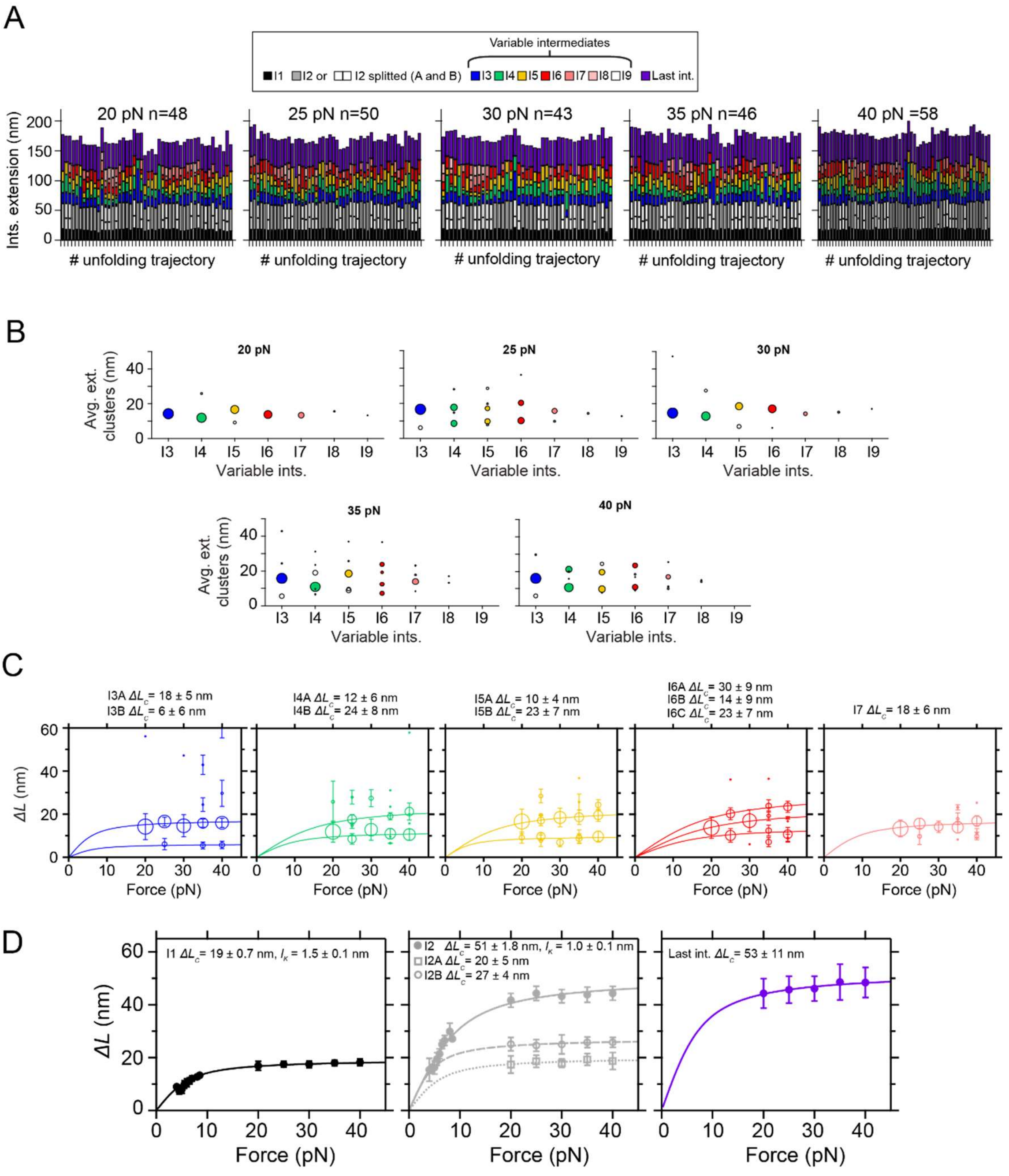


**Supplementary Figure 3. Mechanical characterization of PilY1$^{501\text{-}1161}$ in constant force with $Ca^{2+}$ and identification of the most prevalent conformations of the variable intermediates.** Unfolding under constant force of PilY1$^{501\text{-}1161}$ in the presence of $Ca^{2+}$ and classification of the intermediates by unfolding order and color. The category plots show the stacked extensions of each intermediate per trajectory across the five unfolding forces tested (20, 25, 30, 35, and 40 pN). **B)** Cluster analysis of the extension of the variable intermediates per order of unfolding and force in $Ca^{2+}$. Each plot shows the average extension of the events classified inside each cluster and for each intermediate across the five tested forces. The size of the data points is proportional to the number of events in that cluster (most represented ones are colored). **C)** Force-dependent extensions (mean±SD) of the most populated conformations for each intermediate (I3 to I7). The size of the data points is proportional to the number of events in that cluster. When the extension of the clusters identified in **B** is represented for each intermediate across the five unfolding forces tested, it is possible to identify the different conformations of each intermediate and model the data with individual fits of the FJC model ($\Delta L_c$ data for each population shown above). **D)** Force-dependent extensions (mean±SD) of the I1, I2 (as a single and as two events), and last intermediates and their fitting with the FJC model.

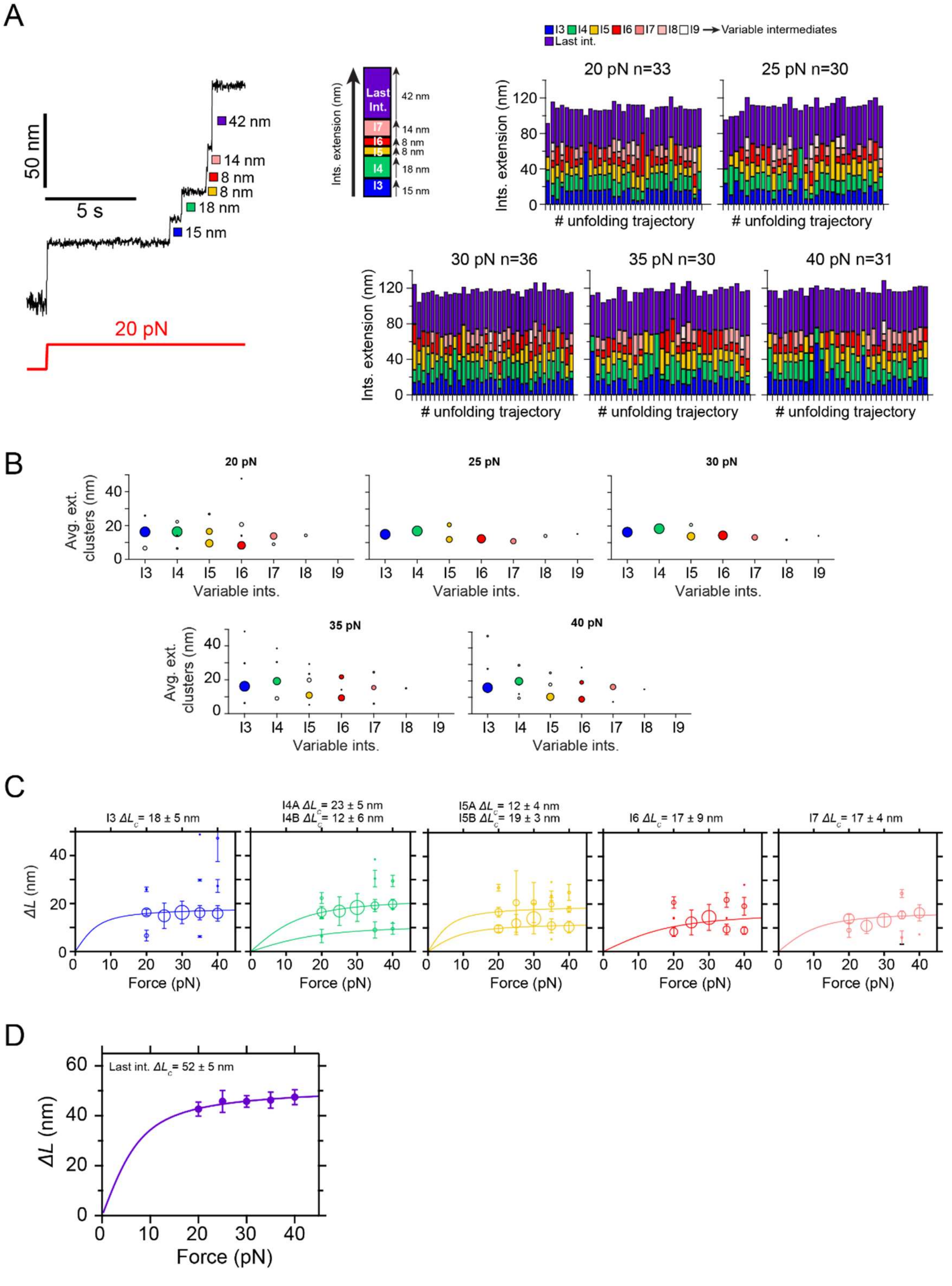


**Supplementary Figure 4. Mechanical characterization of PilY1$^{642\text{-}1161}$ in constant force with EGTA and identification of the most prevalent conformations of the variable intermediates.** Unfolding under constant force of PilY1$^{642\text{-}1161}$ in the presence of EGTA and classification of the intermediates by unfolding order and color. The category plots on the right show the stacked extensions of each intermediate per trajectory across the five unfolding forces tested (20, 25, 30, 35, and 40 pN). This protein lacks the signature of the I1 and I2 intermediates observed in PilY1$^{501\text{-}1161}$. **B)** Cluster analysis of the extension of the variable intermediates per order of unfolding and force in EGTA. Each plot shows the average extension of the events classified inside each cluster and for each intermediate across the five tested forces. The size of the data points is proportional to the number of events in that cluster (most represented ones are colored). **C)** Force-dependent extensions (mean±SD) of the most populated conformations for each intermediate (I3 to I7). The size of the data points is proportional to the number of events in that cluster. When the extension of the clusters identified in **B** is represented for each intermediate across the five unfolding forces tested, it is possible to identify the different conformations of each intermediate and model the data with individual fits of the FJC model ($\Delta L_c$ data for each population shown above). **D)** Force-dependent extension (mean±SD) of the last intermediate and its fitting with the FJC model.

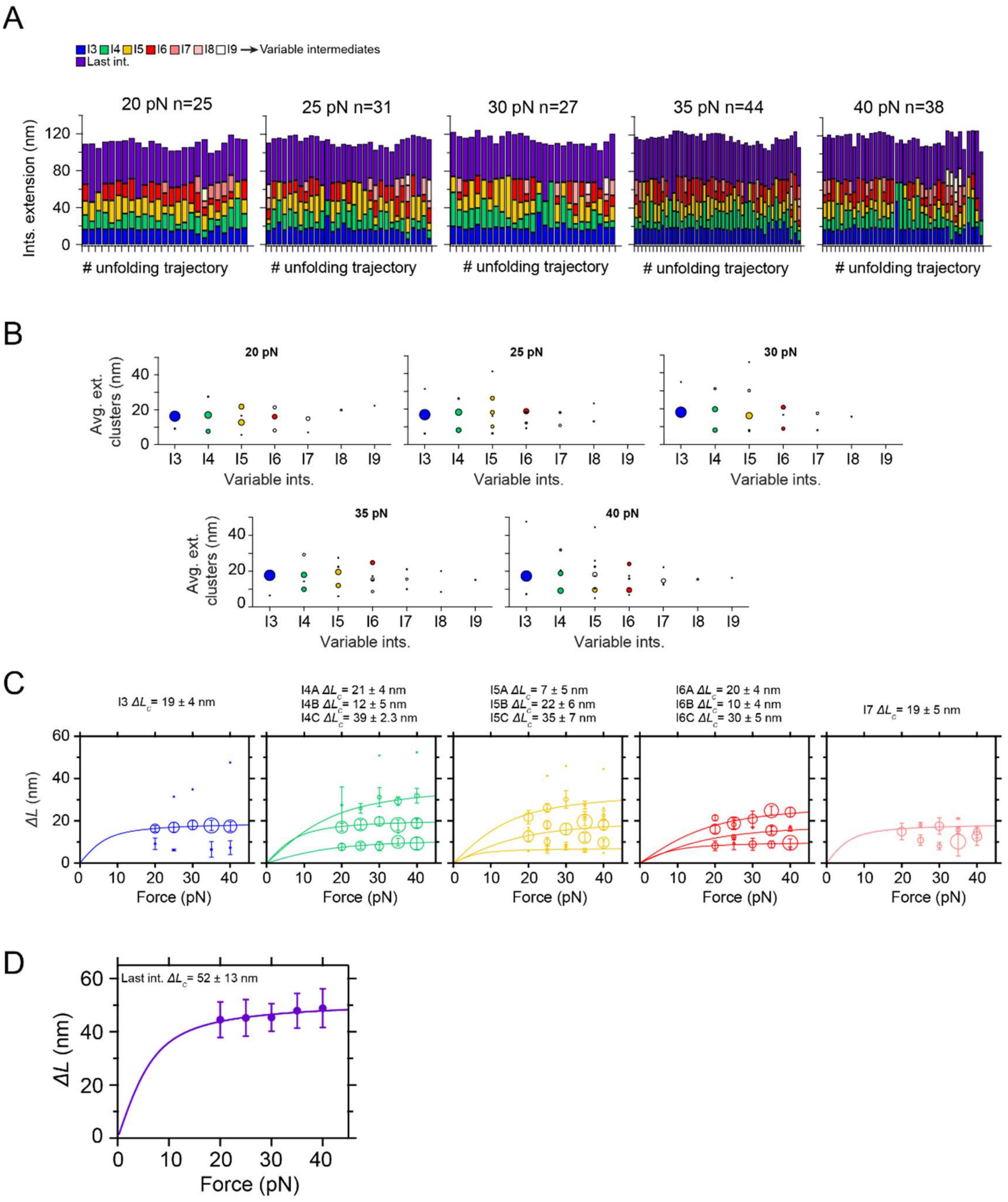


**Supplementary Figure 5. Mechanical characterization of PilY1$^{642\text{-}1161}$ in constant force with $Ca^{2+}$ and identification of the most prevalent conformations of the variable intermediates. A)** Unfolding under constant force of PilY1$^{642\text{-}1161}$ in the presence of $Ca^{2+}$ and classification of the intermediates by unfolding order and color. The category plots show the stacked extensions of each intermediate per trajectory across the five unfolding forces tested (20, 25, 30, 35, and 40 pN). This protein lacks the signature of the I1 and I2 intermediates observed in PilY1$^{501\text{-}1161}$. **B)** Cluster analysis of the extension of the variable intermediates per order of unfolding and force in $Ca^{2+}$. Each plot shows the average extension of the events classified inside each cluster and for each intermediate across the five tested forces. The size of the data points is proportional to the number of events in that cluster (most represented ones are colored). **C)** Force-dependent extensions (mean±SD) of the most populated conformations for each intermediate (I3 to I7). The size of the data points is proportional to the number of events in that cluster. When the extension of the clusters identified in **B** is represented for each intermediate across the five unfolding forces tested, it is possible to identify the different conformations of each intermediate and model the data with individual fits of the FJC model ($\Delta L_c$ data for each population shown above). **D)** Force-dependent extension (mean±SD) of the last intermediate and its fitting with the FJC model.

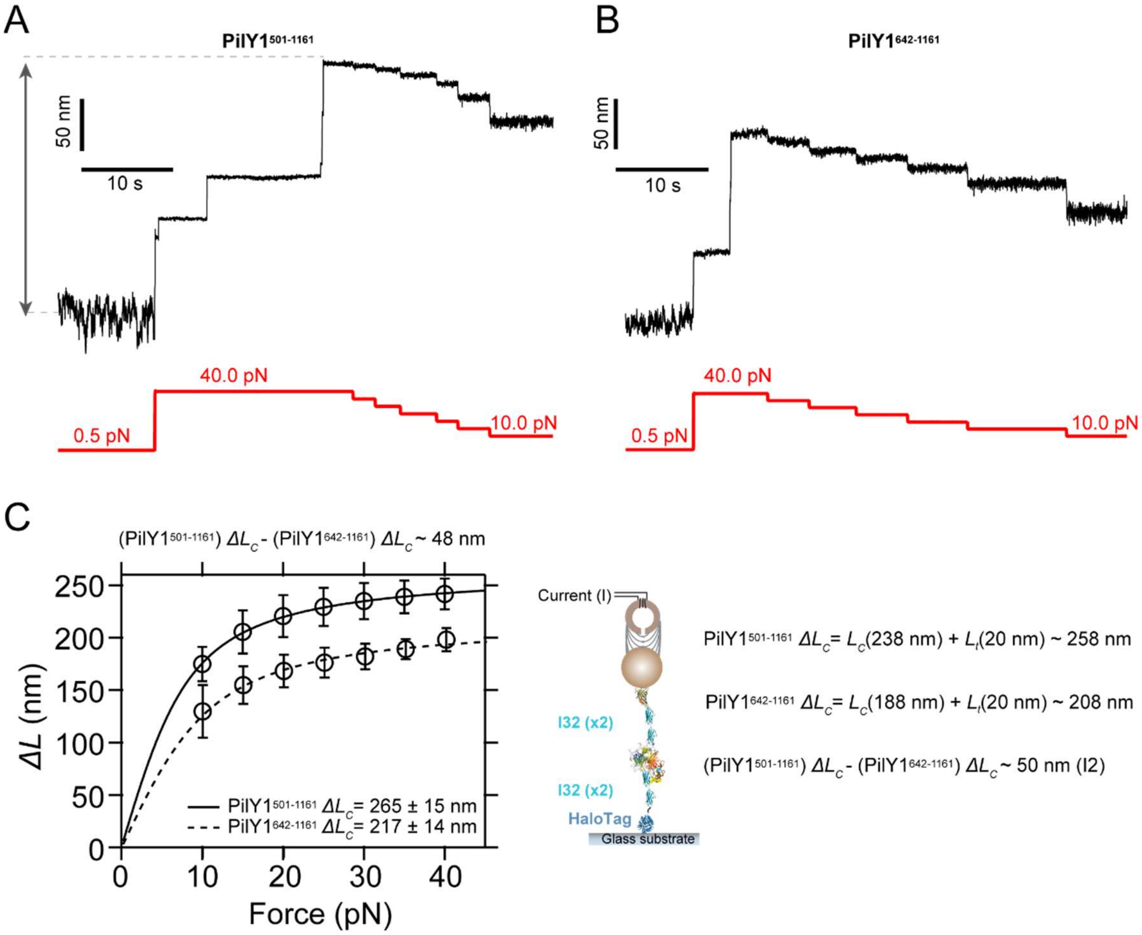


**Supplementary Figure 6. Full extension of PilY1$^{501\text{-}1161}$ and PilY1$^{642\text{-}1161}$. A)** Unfolding extension of PilY1$^{501\text{-}1161}$ from 0.5 pN to 10, 15, 20, 25, 30, 35, and 40 pN. **B)** Unfolding extension of PilY1$^{642\text{-}1161}$ from 0.5 pN to 10, 15, 20, 25, 30, 35, and 40 pN. **C)** Force-dependent extension of both proteins from 0.5 pN to different forces, and fitting to the FJC model. Complete extension ($L_C$) calculated (0.36 nm·res$^{-1}$) using the sequence length and the length of folded domains from the tether after stretching ($L_t$). Both experimentally and theoretically, the difference in length between both constructs is ~50 nm, which matches with the expected size of the I2 intermediate, and which is missing in the construct PilY1$^{642\text{-}1161}$.

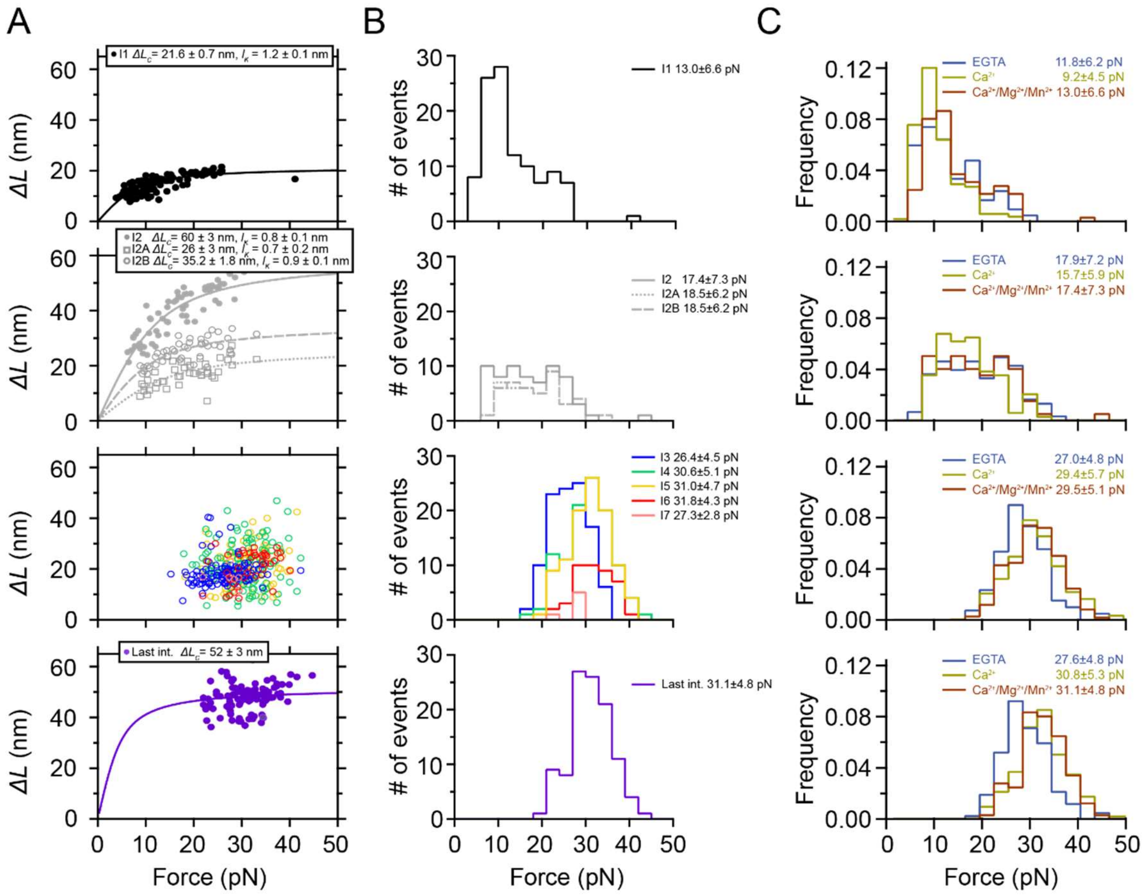


**Supplementary Figure 7. Mechanical characterization of PilY1$^{501\text{-}1161}$ with force-ramp (1 pN·s$^{-1}$) in the presence of binding buffer ($Ca^{2+}$, $Mg^{2+}$, and $Mn^{2+}$). A)** Force-dependent extension of (from top to bottom) I1, I2 (as a single or as two events), variable intermediates, and last intermediate. The lines are fits of the data with the FJC model. **B)** Unfolding force distribution of the four populations of intermediates (mean±SD). Variable intermediates are shown separately. **C)** Comparison of the unfolding force distributions of the four classes of intermediates between the buffer conditions tested. Variable intermediates are shown collectively under a single distribution.

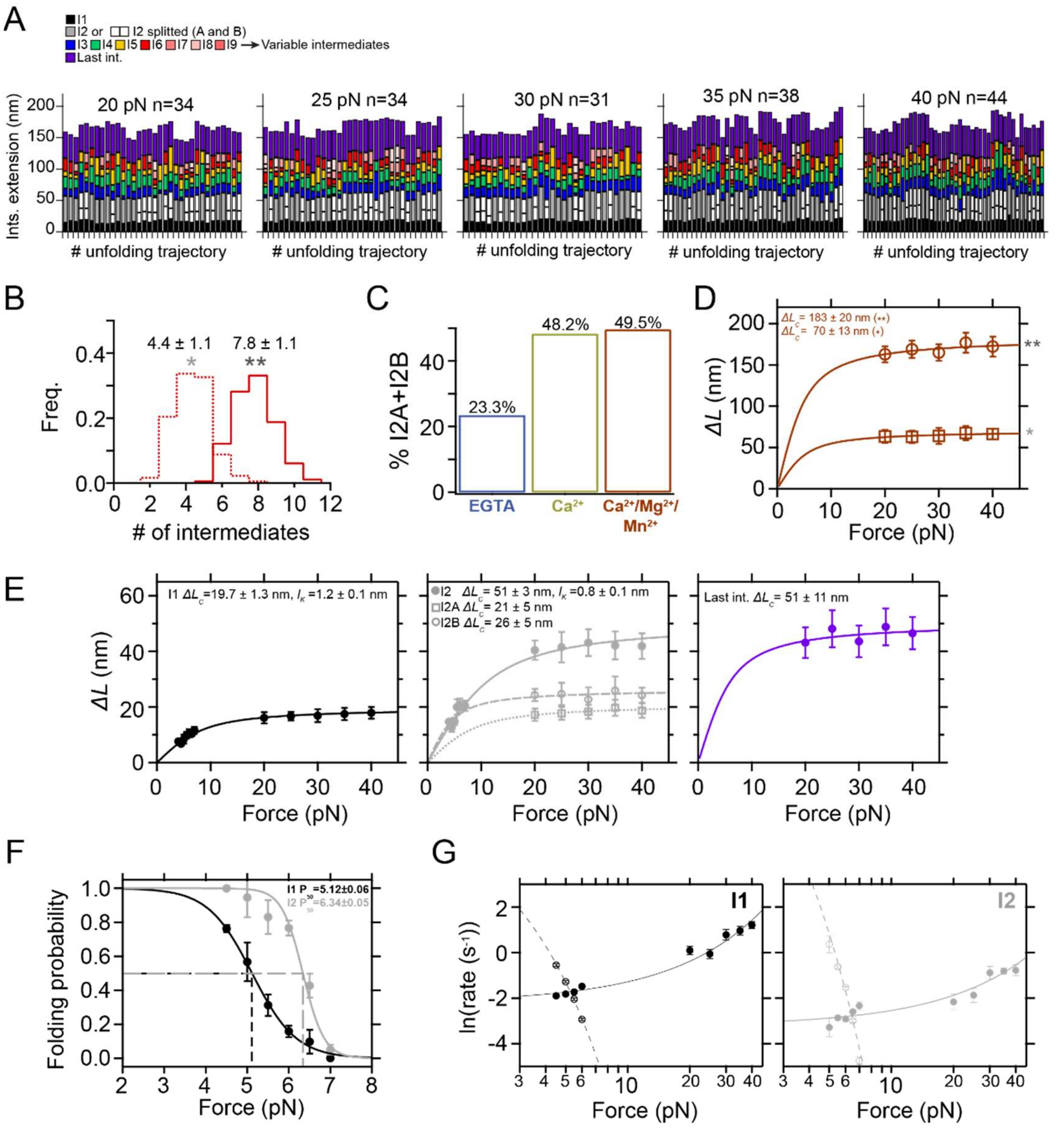


**Supplementary Figure 8. Mechanical characterization of PilY1$^{501\text{-}1161}$ in constant force with binding buffer ($Ca^{2+}$, $Mg^{2+}$, and $Mn^{2+}$). A)** Unfolding under constant force of PilY1$^{501\text{-}1161}$ in binding buffer and classification of the intermediates by unfolding order and color. The category plots show the stacked extensions of each intermediate per trajectory across the five unfolding forces tested (20, 25, 30, 35, and 40 pN). **B)** Distribution of the total number of unfolding intermediates in PilY1$^{501\text{-}1161}$ (**) and variable intermediates (*) observed across trajectories. **C)** Percentage of I2 domains unfolding in two consecutive sub-intermediates compared to EGTA and $Ca^{2+}$ conditions. **D)** Force-dependent change in the combined extension of all PilY1$^{501\text{-}1161}$ intermediates (**) and variable intermediates (*). Lines are fits of the FJC model to the average extension (mean±SD), and the resulting values of contour length increment ($\Delta L_c$) are shown above inside the graph. **E)** Force-dependent extension change (mean±SD) of the I1, I2 (as a single and as two events), and last intermediates and their fitting with the FJC model. **F)** Folding probability of I1 and I2 between 4.5 and 7.0 pN. The folded fraction of both intermediates sharply decreases with force, following a sigmoidal trend (line fits). The coexistence force ($P_{50}$) of I1 is 5.1 pN, and 6.3 pN for I2. **G)** Force-dependency of folding and unfolding kinetics of I1 and I2. I1 and I2 unfolding and folding kinetics with Bell's model (**I1**, empty circles and dashed line for folding: $k_F^0 = (8.0 \pm 2.1) \times 10^2$ s$^{-1}$, $\Delta x_F = -6.56 \pm 0.20$ nm. Solid circles and lines for unfolding: $k_U^0 = (11.2 \pm 0.04) \times 10^{-2}$ s$^{-1}$, $\Delta x_U = 0.37 \pm 0.01$ nm; **I2** $k_F^0 = (7.2 \pm 3.6) \times 10^5$ s$^{-1}$, $\Delta x_F = -10.41 \pm 0.34$ nm. Solid circles and lines for unfolding: $k_U^0 = (4.0 \pm 0.2) \times 10^{-2}$ s$^{-1}$, $\Delta x_U = 0.27 \pm 0.01$ nm).

| DOMAINS | CANDIDATE INTERMEDIATES |
|---|---|
| IBD[501-643] $L_0$ = 1.0 nm, $\Delta L_c$= 50 nm 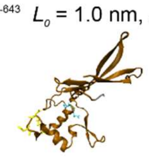 | I2 $\Delta L_c$= 51 ± 1.8 nm |
| Blade I[642-716] $L_0$ = 2.9 nm, $\Delta L_c$= 23.7 nm 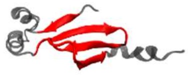 | I1 $\Delta L_c$= 19 ± 0.7 nm<br>I3 $\Delta L_c$= 18 ± 5 nm<br>I5B $\Delta L_c$= 23 ± 7 nm<br>I6C $\Delta L_c$= 23 ± 7 nm |
| Blade II[717-783] $L_0$ = 2.8 nm, $\Delta L_c$= 21 nm 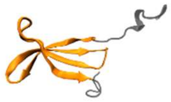 | I1 $\Delta L_c$= 19 ± 0.7 nm<br>I3A $\Delta L_c$= 18 ± 5 nm<br>I5B $\Delta L_c$= 23 ± 7 nm<br>I6C $\Delta L_c$= 23 ± 7 nm |
| Blade III[783-838] $L_0$ = 1.5 nm, $\Delta L_c$= 18.3 nm 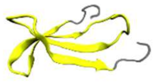 | I1 $\Delta L_c$= 19 ± 0.7 nm<br>I3A $\Delta L_c$= 18 ± 5 nm<br>I5B $\Delta L_c$= 23 ± 7 nm<br>I6C $\Delta L_c$= 23 ± 7 nm |
| Blade IV[838-916] $L_0$ = 1.1 nm, $\Delta L_c$= 27 nm 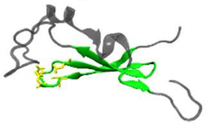 | I4B $\Delta L_c$= 24 ± 8 nm<br>I5B $\Delta L_c$= 23 ± 7 nm<br>I6C $\Delta L_c$= 23 ± 7 nm |
| Blade V-VI[916-1119] $L_0$ = 2.5 nm, $\Delta L_c$= 70.6 nm 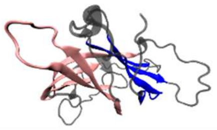 | Last int. $\Delta L_c$= 53 ± 11 nm |
| Blade VII[1119-1163] $L_0$ = 5.2 nm, $\Delta L_c$= 10.6 nm 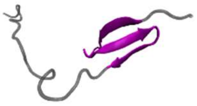 | I4A $\Delta L_c$= 12 ± 6 nm<br>I5A $\Delta L_c$= 10 ± 4 nm<br>I6B $\Delta L_c$= 14 ± 9 nm |

**Supplementary Figure 9. List of PilY1[501-1161] domains and a rough estimation of their size and candidate intermediates.** On the left, a cartoon representation of each domain. The domains are connected by α-helices and unstructured loops, and the criteria for defining the ends of each domain (N-C distance in the folded structure, $L_0$) has been set by splitting by half the length of these interdomain regions. Contour length values ($L_C$) are determined by multiplying the number of residues of each module by 0.36 $nm \cdot res^{-1}$. On the right, $\Delta L_C$ values for each domain of PilY1[501-1161] in $Ca^{2+}$ (**Supplementary Figure 3**). Due to their overlapping sizes, most of the β-propeller blades have several candidates. Blades V-VI have been represented as a single unit because they share a β-strand that splits both domains. Hence, their structural integrity could be interdependent. Under force, the interdomain regions could be stretched before unfolding; therefore, the estimated values represented here could be overestimated.
